# High throughput method for simultaneous screening of membrane permeability and toxicity for discovery of new cryoprotective agents

**DOI:** 10.1101/2024.07.22.604685

**Authors:** Nima Ahmadkhani, James D. Benson, Ali Eroglu, Adam Z. Higgins

## Abstract

Vitrification is the most promising method for cryopreservation of complex structures such as organs and tissue constructs. However, this method requires multimolar concentrations of cell-permeant cryoprotective agents (CPAs), which can be toxic at such elevated levels. The selection of CPAs for organ vitrification has been limited to a few chemicals; however, there are numerous chemicals with properties similar to commonly used CPAs. In this study, we developed a high-throughput method that significantly increases the speed of cell membrane permeability measurement, enabling ~100 times faster permeability measurement than previous methods. The method also allows assessment of CPA toxicity using the same 96-well plate. We tested five commonly used CPAs and 22 less common ones at both 4 °C and room temperature, with 23 of them passing the screening process based on their favorable toxicity and permeability properties. Considering its advantages such as high throughput measurement of membrane permeability along with simultaneous toxicity assessment, the presented method holds promise as an effective initial screening tool to identify new CPAs for cryopreservation.

**Significance:** Cryoprotective agent (CPA) toxicity is the most limiting factor impeding cryopreservation of critically needed tissues and organs for transplantation and medical research. This limitation is in part due to the challenge of rapidly screening compounds to identify candidate molecules that are highly membrane permeable and non-toxic at high concentrations. Such a combination would facilitate rapid CPA permeation throughout the sample, enabling ice-free cryopreservation with minimal toxicity. This study presents a method for rapidly assessing the cell membrane permeability and toxicity of candidate CPAs, identifies several novel high-permeability low-toxicity CPAs for further testing, and lays the groundwork for additional high throughput screening to discover novel CPAs with the potential to improve cryopreservation of complex tissues and organs.

## 1. Introduction

Reproducible cryopreservation of spermatozoa was achieved upon discovery of the cryoprotective properties of glycerol in 1949 [1] and it is now common to cryopreserve isolated cells including spermatozoa, oocytes, and cell lines. Nevertheless, progress in the cryopreservation of complex biological samples such as tissues and organs has been relatively limited [2–4]. Over 100,000 people in the United States are currently on organ transplantation waiting lists, and thousands die each year because there are not enough organs [5]. The organ shortage problem is further compounded by the issue of rejection after transplantation [6]. Current hypothermic organ preservation methods provide short preservation times, ranging from 6 h for heart to 24-36 h for kidneys [7]. This can lead to the unfortunate discarding of a significant portion of donated organs [8]. Moreover, the inability to conduct comprehensive immunological assessments due to these time constraints contributes to organ rejection [6, 8]. Cryopreservation would provide researchers with ample time to conduct comprehensive studies on the organs, enabling them to better understand the reasons for rejection and work towards effective solutions [8].

The advantages of cryopreservation extend beyond organs. Cryopreservation is becoming increasingly crucial in regenerative medicine for ensuring the provision of secure and standardized products for both patient treatments and research laboratories [9]. To achieve this objective, effective cryopreservation of adherent cells is imperative [10, 11], including cells within 3-dimensional hydrogels [12, 13], cell sheets [14], and cell-based biosensor chips [15, 16]. Cryopreservation is essential for the long-term storage and transportation of these products to their final destinations [17].

There are two primary methods of cryopreservation: slow freezing and vitrification. Vitrification, in particular, holds promise because it prevents ice formation, thereby eliminating the associated damage [18]. Vitrification requires the use of multimolar concentrations of chemicals known as cryoprotectants or cryoprotective agents (CPAs). These chemicals should ideally permeate the entire sample, including the intracellular space, to suppress ice formation. One of the major obstacles to cryopreservation of complex samples is identification of CPA mixtures that prevent ice formation without killing the cells due to toxicity [4]. This is particularly true for bulky samples like organs, because even higher CPA concentrations are needed to prevent ice formation at the relatively slow cooling and warming rates achievable in large samples. Despite these formidable challenges, there is evidence that cryopreservation of complex samples like organs may be achievable, including a report published in 2009 describing survival of a single cryopreserved rabbit kidney after transplantation [19], and a report published just last year demonstrating survival and life-sustaining function after transplantation of cryopreserved rat kidneys [2]. In both cases, the surviving kidneys showed signs of damage, most likely because of CPA toxicity or perfusion injury. Moreover, the CPA concentrations used in these studies may not be sufficient to prevent ice formation in larger human organs. This highlights the need for new, less toxic, CPA mixtures.

Previous studies of tissue and organ cryopreservation have almost entirely been limited to a small subset of the possible chemicals that could be used as CPAs. The key properties shared by commonly used CPAs are low toxicity, the ability to penetrate cell membranes, and the ability to inhibit ice formation in aqueous mixtures. There are many chemicals that may have favorable properties that have not been tested for cryopreservation. To identify promising leads, high throughput screening tools are needed, including methods to screen for chemicals that are membrane permeable and relatively nontoxic.

Various methods have been used to measure CPA permeability, such as microfluidic systems [20–22], Coulter counters [23] and microscopic measurement of fluorescence quenching [24]. Most of these methods are applicable to suspended cells and require more than 15 minutes for a single permeability measurement. The only method applicable to adherent cells is a fluorescence quenching technique that was previously conducted in our lab; however, it requires about 30 minutes per sample [24]. We previously demonstrated a high-throughput method for toxicity screening in plated cells [25], but to our knowledge, there is currently no high-throughput method available for measuring cell membrane permeability.

In this study, we adapted the fluorescence quenching method for use with an automated plate reader, enabling permeability measurements for an entire 96-well plate in about 30 minutes. The same well plate was then used to estimate CPA toxicity. This method was used to assess the permeability and toxicity of 27 chemicals at two different temperatures, 4 °C and room temperature. Our results demonstrate the potential of this method as an initial screening tool for the discovery of new chemicals with promising permeability and toxicity characteristics, and for development of more effective CPA mixtures for cryopreservation.

## 2. Results

To address the need for discovery of more effective CPAs for cryopreservation, we developed a new method for high throughput screening of candidate chemicals to assess cell membrane permeability and toxicity using an automated plate reader (Figure 1). This method was applied to bovine pulmonary artery endothelial cells cultured in 96-well plates for 27 test chemicals (plus sucrose as a non-permeant control) at 4 °C and 25 °C.

**Figure 1.**
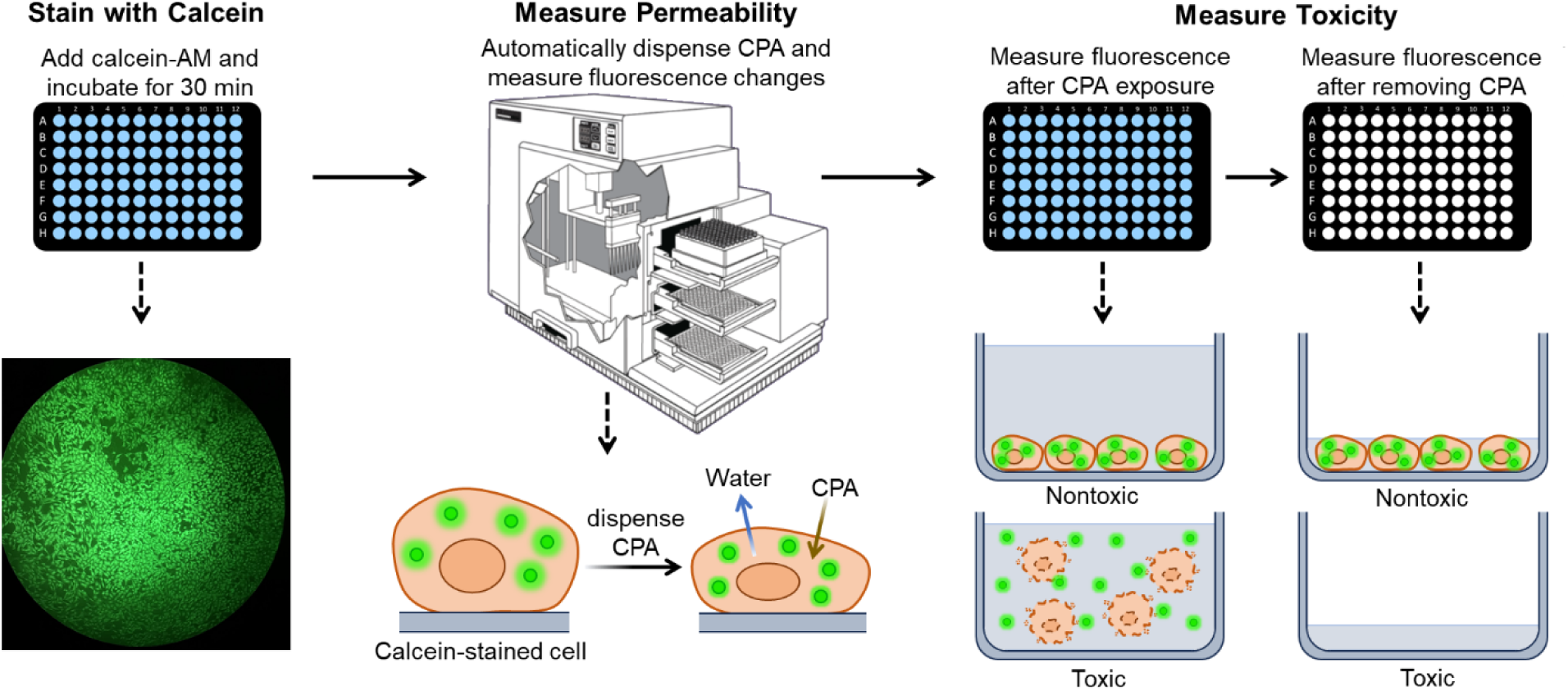
High throughput method for measuring cell membrane permeability and toxicity. Calcein-stained cells in 96-well plates were exposed to candidate CPAs and the resulting changes in fluorescence were used to estimate the cell membrane permeability. The same plate was then used to estimate CPA toxicity by comparing the fluorescence before and after removal of the solution from the wells. The image of the FlexStation 3 plate reader was used with permission from Molecular Devices LLC (https://www.moleculardevices.com).

### 2.1. Cell Membrane Permeability

The method for measuring cell membrane permeability is based on the use of intracellular calcein as a marker of cell volume. Calcein fluorescence has previously been shown to be quenched by molecules present in the cell cytoplasm, resulting in an approximately linear relationship between fluorescence and cell volume [24, 26]. Using the automated plate reader, we observed similar changes in relative fluorescence intensity after inducing cell volume changes by exposing the cells to solutions containing nonpermeating solutes. Hypertonic solutions cause cell water volume loss, while hypotonic solutions cause cell water volume gain. Likewise, cell water volume loss results in a decrease in fluorescence, whereas cell water volume gain results in an increase, as illustrated in Figure 2A. Figure 2B shows the equilibrium normalized cell fluorescence *F̄* for the isotonic condition and three different hypertonic solution concentrations. The Boyle-van’t Hoff equation (Eq. 5) was used to express the relative concentration on the horizontal axis in terms of the relative osmotically active cell volume *V̄*. Thus, Figure 2B can be used to examine the relationship between *F̄* and *V̄*. As depicted in Figure 2B, there is a linear relationship between normalized cell fluorescence and normalized osmotically active cell volume when cells are exposed to hypertonic solutions (R^2^ = 0.99). Figure 2B focuses on hypertonic conditions that cause cell shrinkage because our goal is to measure CPA permeability and exposure to hyperosmotic CPA solutions also causes shrinkage. In contrast, the equilibrium fluorescence under hypotonic conditions (which causes swelling) deviates from the linear trend (Figure S3).

**Figure 2.**
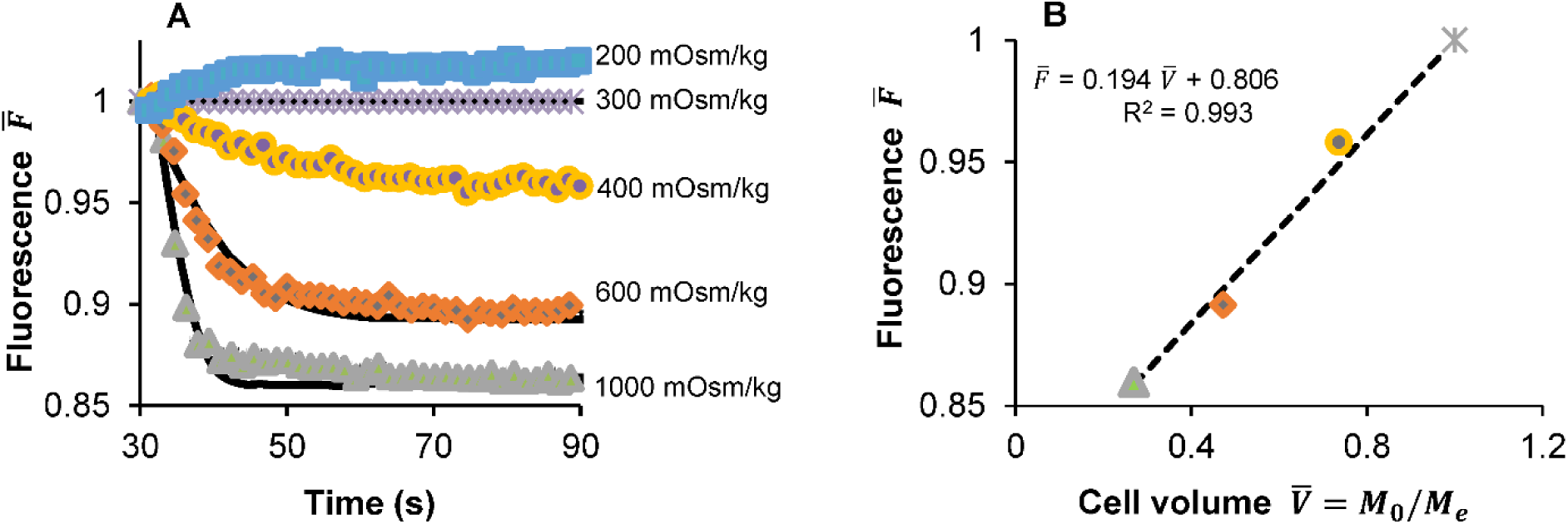
Relationship between fluorescence and expected cell volume. A) Change in normalized fluorescence over time for different concentrations of hypertonic and hypotonic solutions containing non-permeating solutes. B) Equilibrium fluorescence as a function of predicted equilibrium cell volume (see Eq. 5) after exposure to hypertonic solutions at various concentrations. Each data point represents the average ± SEM for 10-12 wells. Experiments were performed at 25 °C.

Figure 3 illustrates the patterns observed in fluorescence intensity when cells were exposed to 28 different chemicals at 4 °C. The impact of the nonpermeating solute sucrose on cell fluorescence is demonstrated in the upper left panel. Exposure to sucrose causes cell water loss that plateaus at a stable volume at a low level of fluorescence. As expected, the best-fit solute permeability for sucrose was very low (*P̄_CPA_* = 0.00026 ± 0.00007). Exposure to a permeating solute typically causes initial cell shrinkage followed by a slower increase in cell volume as the solute permeates the cell membrane. This is illustrated by the panels with yellow diamonds, which are presented in order of ascending solute permeability. All these chemicals had a solute permeability that was significantly higher than that of sucrose. The panels with blue circles exhibit a minor rise in fluorescence, accompanied by minimal to no indication of cell shrinkage. This is consistent with rapid solute permeation. In these cases, it was not possible to fit the data to determine the membrane permeability parameters. The remaining panels (red triangles) show the results for chemicals that were found to be toxic.

**Figure 3.**
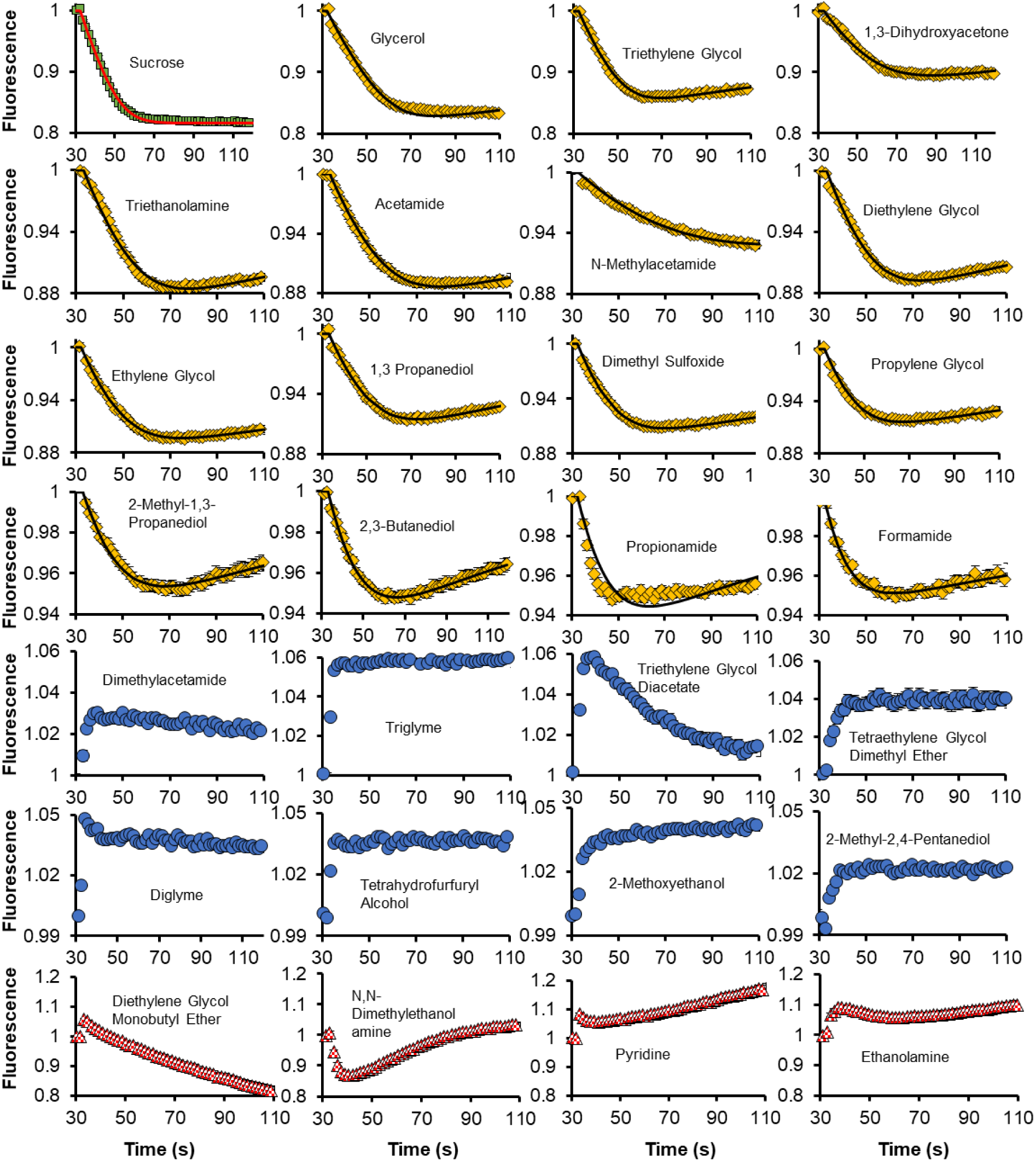
Transient fluorescence data (symbols) and permeability model predictions (lines) after exposure to various chemicals at 4 °C. The solutions had a total osmolality of 1000 ± 100 mOsm/kg. Each data point represents the average ± SEM of a minimum of 13 replicates conducted across at least 3 plates.

Similar trends were observed at 25 °C, as shown in Figure 4. The main difference is that the fluorescence response was faster at 25 °C than at 4 °C, and more of the candidate CPAs were found to be toxic. Table 1 summarizes the best-fit permeability parameters at 4 °C and 25 °C. Several chemicals in this table exhibit exceptionally fast membrane permeation, exceeding our measurement capabilities. These chemicals belong to the category of “fast” membrane-permeable CPAs. Table 2 presents the calculated activation energy values for the transport of water and CPA. The activation energies were determined using Arrhenius models (Eqs. 6 and 7).

**Figure 4.**
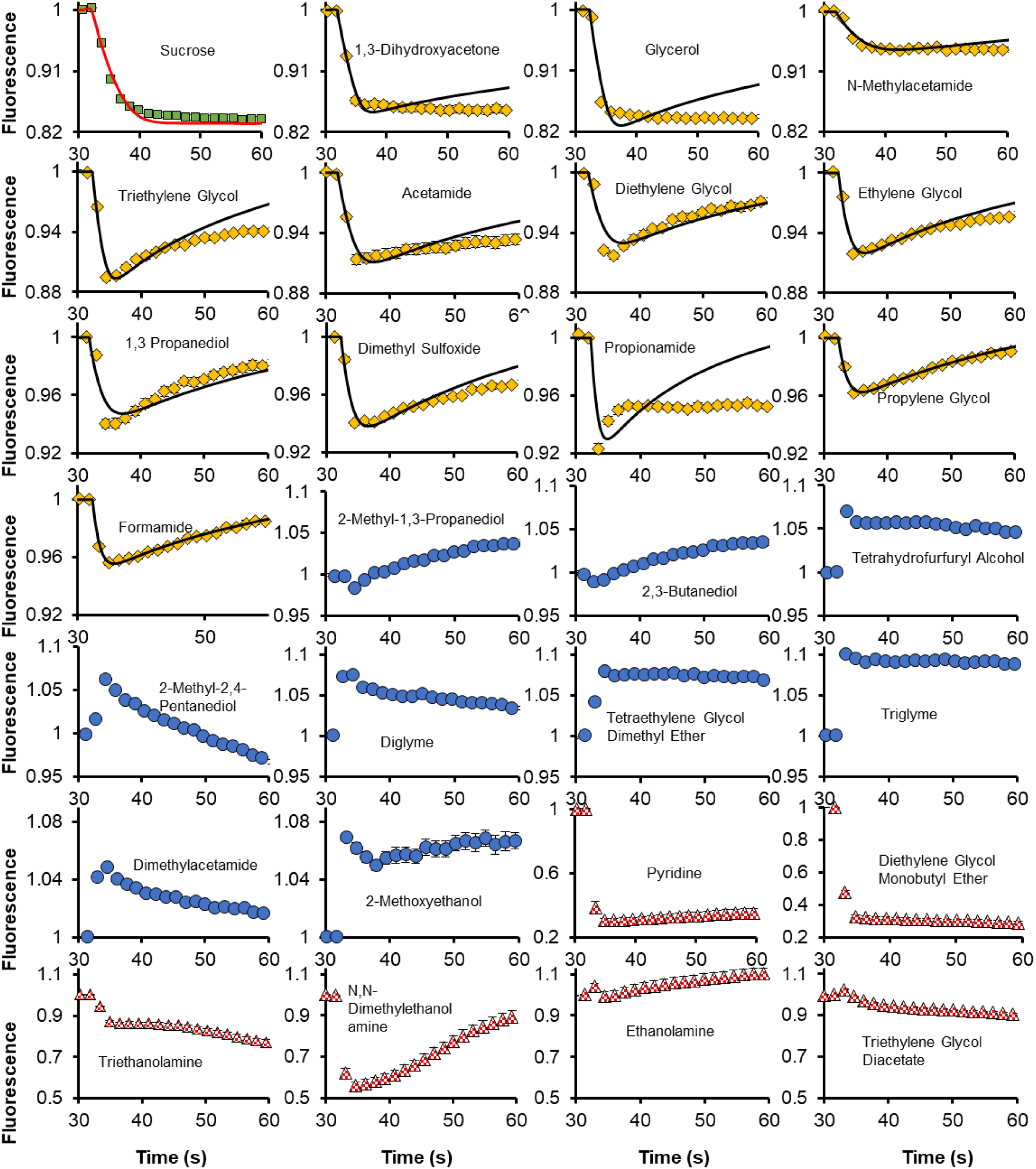
Transient fluorescence data (symbols) and permeability model predictions (lines) after exposure to various solutes at 25 °C. The solutions had a total osmolality of 2000 ± 100 mOsm/kg, except for sucrose, which had an osmolality of 1000 ± 100 mOsm/kg. Each data point represents the average ± SEM of a minimum of 16 replicates conducted across at least 3 plates.

**Table 1.**
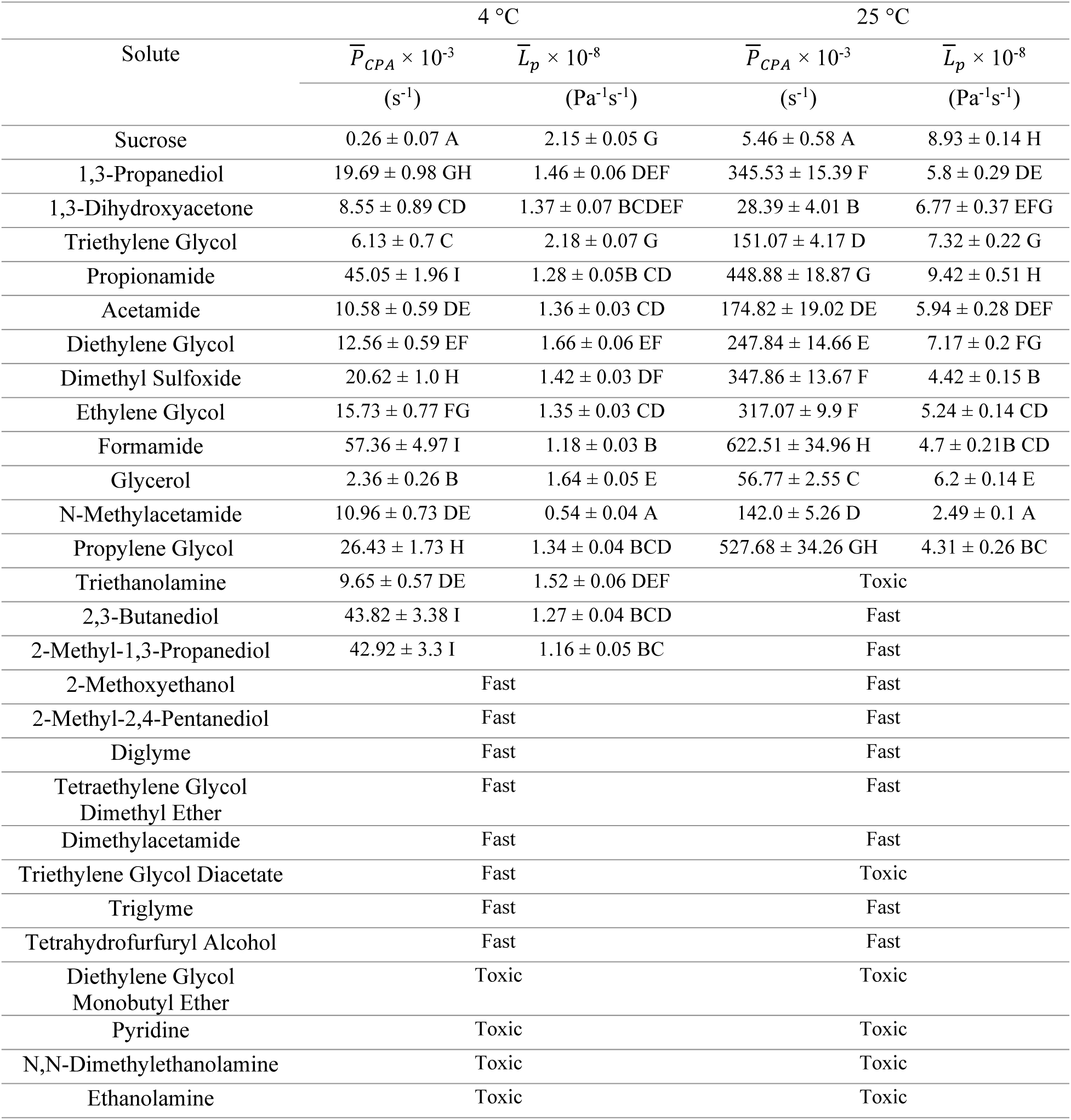
Comparisons of permeability parameters for different chemicals at 4 °C and 25 °C. At the given temperatures, statistical differences in permeability values are indicated by distinct letters. Fast: fluorescence response consistent with solute permeation that was too fast to measure. Toxic: the solute was found to be toxic, so permeability was not measured.

**Table 2.**
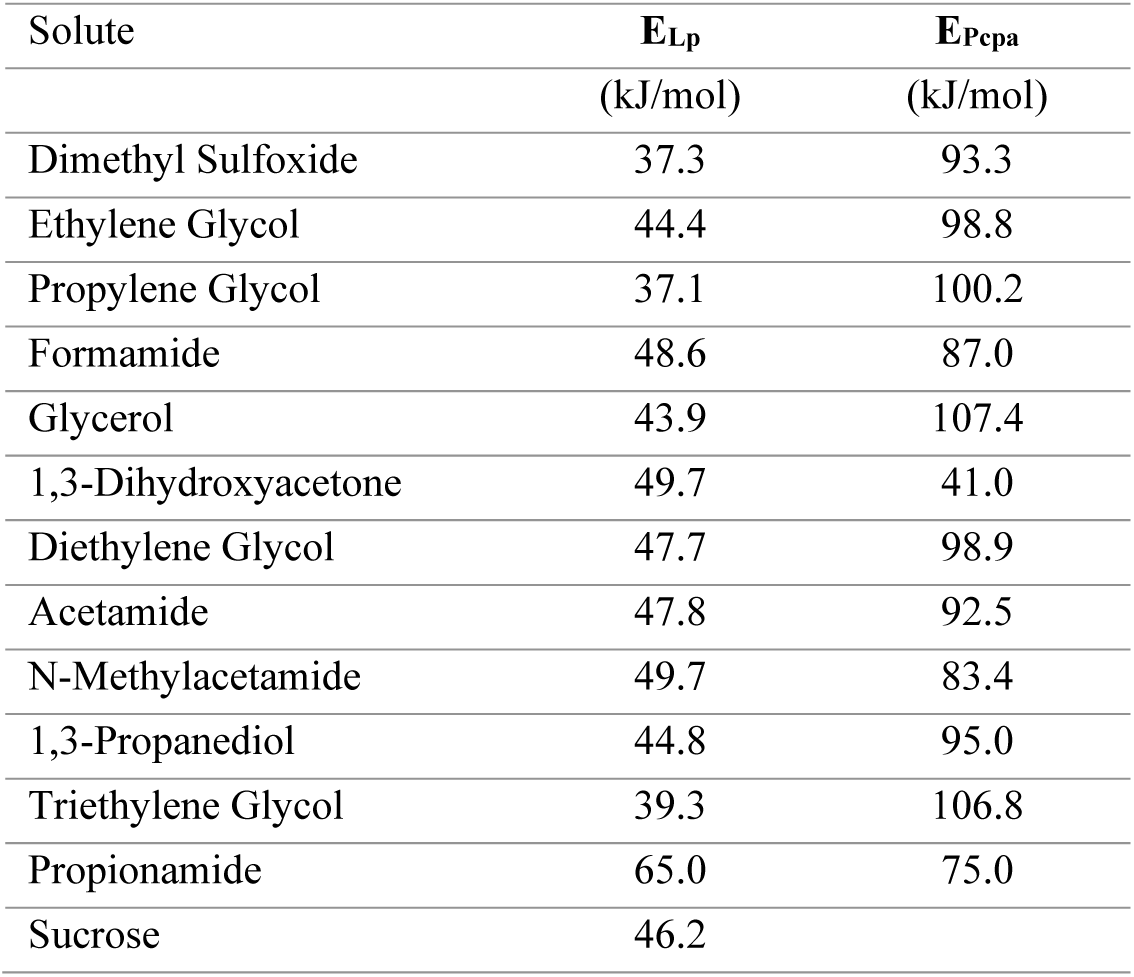
Activation energies for water and solute transport.

### 2.2. CPA toxicity

After measuring the cell membrane permeability parameters, the same well plate was used to assess the toxicity of the candidate CPAs. To perform the toxicity assay, fluorescence was measured after ~20 min CPA exposure, and then again after removing the CPA solution from the wells (see Figure 1). Our strategy for assessing viability assumes that healthy cells retain intracellular calcein, while dead cells with compromised membranes release calcein into the surrounding solution. To examine this assumption, we collected microscopic images after exposure to various candidate CPAs. Figure 5 shows representative images for a relatively nontoxic chemical that was found to yield high viability (left) and a toxic chemical that yielded low viability (right). For nontoxic chemicals, background fluorescence is minimal, and the cells exhibit relatively high fluorescence. In contrast, in wells exposed to toxic chemicals, background fluorescence is high, consistent with loss of intracellular calcein to the surrounding medium.

**Figure 5.**
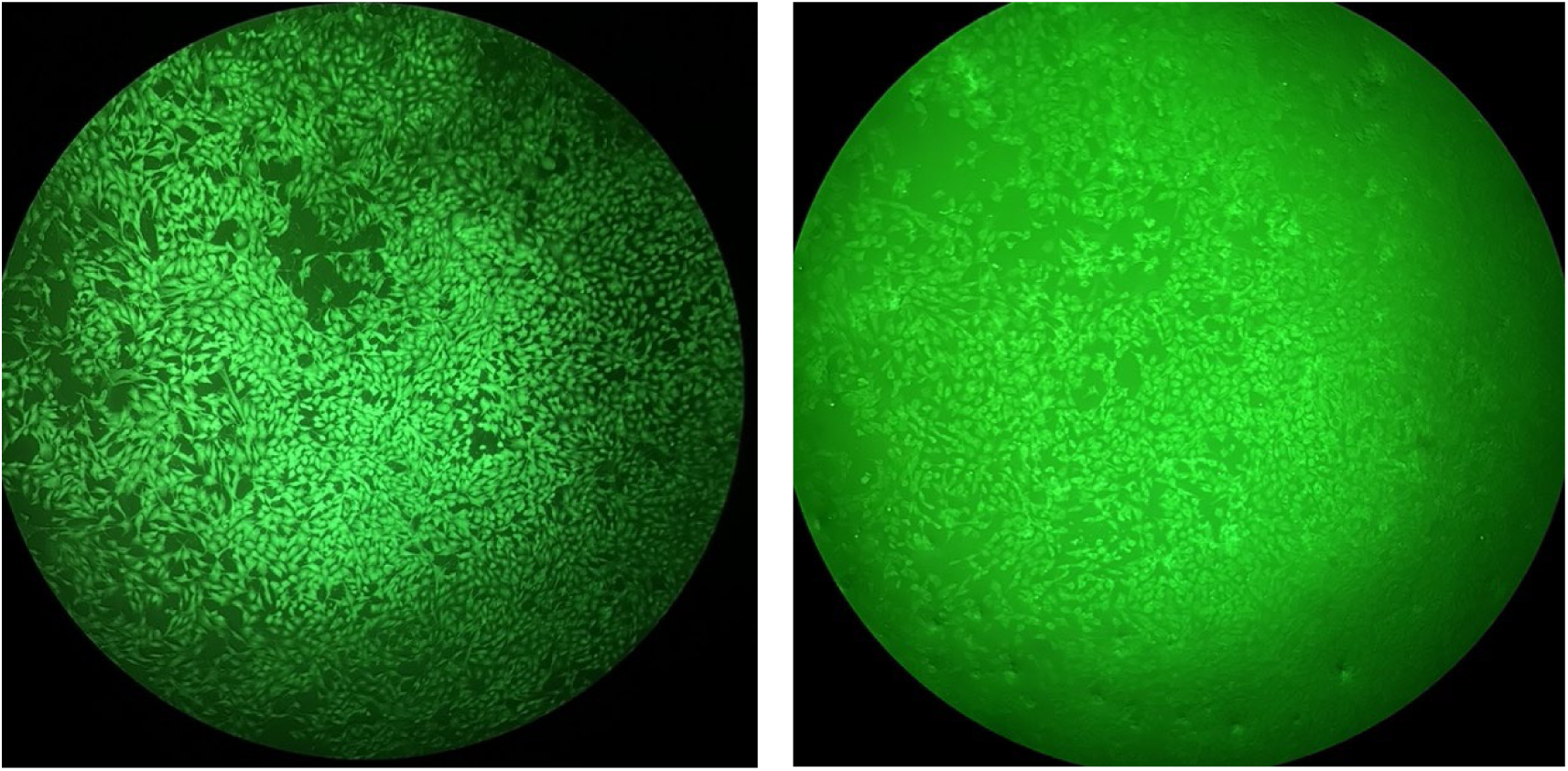
Representative images of calcein-stained cells after exposure to a relatively nontoxic chemical (diglyme, left), and a relatively toxic chemical (ethanolamine, right).

Figure 6A illustrates cell viability measurements plotted against CPA permeability for 27 different candidate CPAs at 4 °C. We used a threshold of 80% for distinguishing between toxic and non-toxic chemicals. Four chemicals were found to be toxic: N,N-dimethylethanolamine, ethanolamine, diethylene glycol monobutyl ether, and pyridine. The permeability of the toxic chemicals, located on the left side of the graph, could not be measured. Figure 6A facilitates identification of chemicals with potential to be effective CPAs. It is desirable for CPAs to have low toxicity and high permeability, which corresponds to the upper right of the graph.

**Figure 6.**
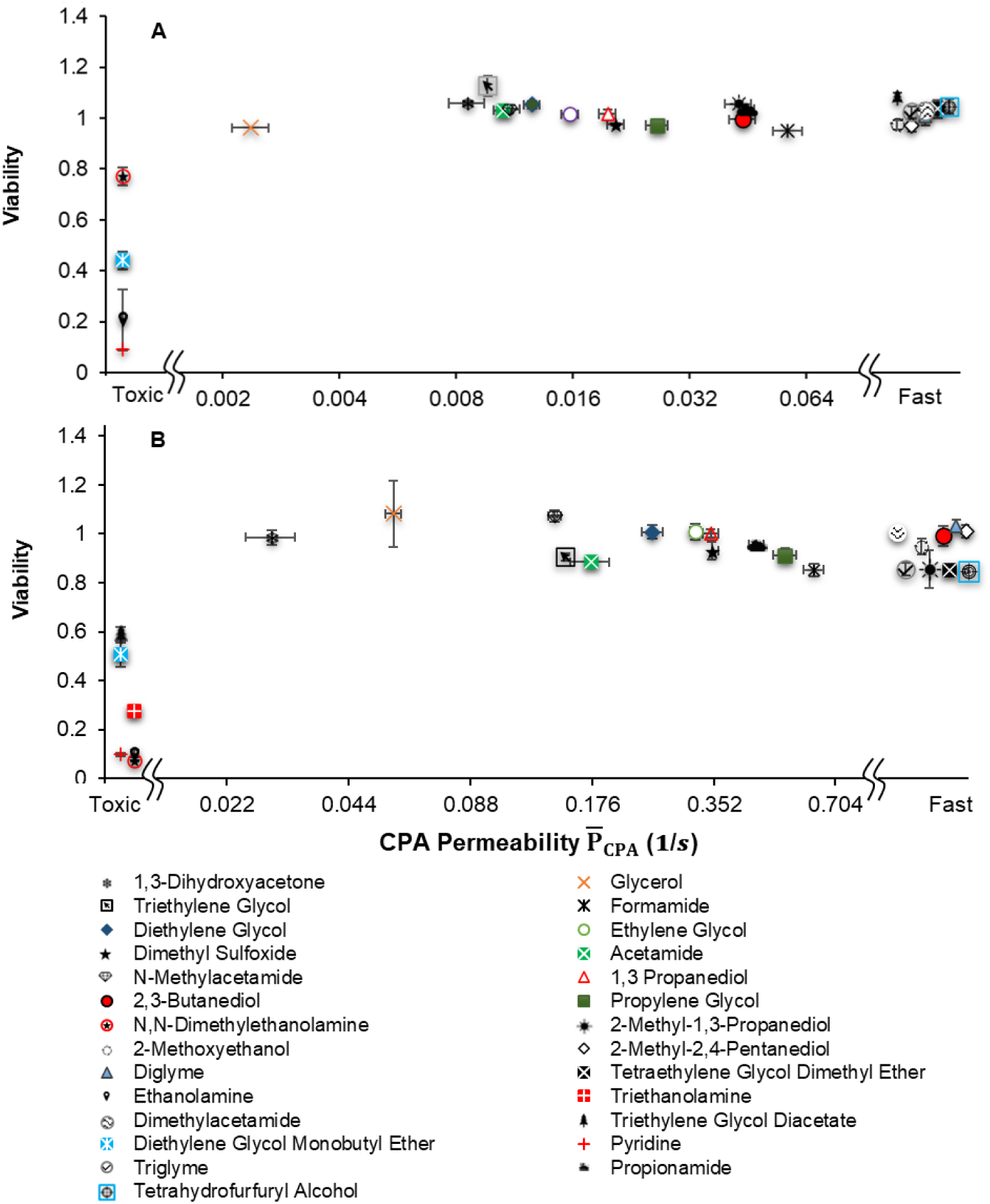
Cell viability and solute permeability. A) Cell viability after ~25 min exposure to ~1 molar solutions at 4 °C as a function of solute permeability. Each data point represents the average ± SEM of a minimum of 13 permeability replicates and 4 viability replicates. B) Cell viability after ~20 min exposure to ~ 2 molar solutions at 25 °C as a function of solute permeability. Each data point represents the average ± SEM of a minimum of 16 permeability replicates and 4 viability replicates.

Figure 6B depicts the cell viability and permeability observed after subjecting the cells to the 27 candidate CPAs at a temperature of 25°C. In this case, six chemicals were found to be toxic: triethanolamine, triethylene glycol diacetate, N,N-dimethylethanolamine, ethanolamine, diethylene glycol monobutyl ether, and pyridine.

## 3. Discussion

In this study, we have devised a high-throughput method to assess cell membrane permeability and toxicity for adherent cells in a 96-well plate. Unlike many conventional methods for measuring cell membrane permeability, which are typically performed on cell suspensions (e.g., Coulter counter method [11, 27]), this approach not only provides rapid and consistent results but also in situ measurement of membrane permeability in adherent cells. In our earlier research, we quantified the permeability of adherent cells by employing the calcein fluorescence quenching method in combination with a perfusion chamber and microscopic imaging. A significant limitation of this approach was that it still required one-by-one sample measurement, similar to other conventional methods [24]. Previous methods, including calcein fluorescence quenching and Coulter counter methods, require 15-30 minutes for a single permeability measurement [23, 24]. This is significantly slower compared to our new method, which can measure the permeability for approximately 100 samples in just around 30 minutes, while also enabling estimation of CPA toxicity.

The high throughput method reported here opens new possibilities for discovery of chemicals with promising properties for cryopreservation. To be effective, CPAs should ideally permeate cell membranes and prevent ice formation, without causing toxicity. Our new method can be used to screen for chemicals with high membrane permeability and low toxicity. We used 80% viability as a threshold to screen for chemicals with low toxicity. While there is not a clear threshold for CPA permeability, it is well known that chemicals with relatively low permeability like glycerol increase the risk of osmotic damage. Thus, the upper right region of Figure 6 highlights CPAs with promising permeability properties and low toxicity. As shown in Figure 6A, 23 chemicals were nontoxic during short term (~25 min) exposure to ~1 molar CPA at 4 °C, all of which had CPA permeability values greater than or equal to the permeability of glycerol. Only five of these chemicals are commonly used for cryopreservation. This underscores the need for additional studies to further characterize the effectiveness of these chemicals for cryopreservation.

Overall, our permeability results are consistent with the expected trends based on previous studies. Compared to our previous study, which also assessed the permeability of bovine pulmonary artery endothelial cells, the CPA permeability trend remained consistent (i.e., glycerol had the lowest permeability, followed by ethylene glycol, DMSO, and propylene glycol [24]). However, the CPA permeability values measured in the current study were about 2 times higher at 4 °C and about 4 times higher room temperature (Figure S11). One possible reason for these differences is that the cells originated from different source vials, and thus likely came from different animals. While most previous studies are limited to common CPAs, Naccache and Sha’afi [28] measured the permeability of a wide range of chemicals in human red blood cells, 17 of which overlapped with the chemicals tested in the current study. The permeability trend for these 17 chemicals was similar even though two different cell types were tested. As shown in Figure S12, the permeability values from these studies exhibited an approximately linear trend at both 4 °C (R^2^ = 0.80) and room temperature (R^2^ = 0.71). Overall, the magnitude of the permeability values determined in this study are consistent with the literature [29] and our data are consistent with the common observation that glycerol has a particularly low membrane permeability [23, 30]. As expected, we observed lower permeability values at lower temperature, and the activation energies for water and CPA transport determined in this study are consistent with previous studies [23, 24, 31–33].

There are certain limitations associated with the developed method, which are inherent to all techniques utilizing calcein fluorescence quenching. The calcein method is limited because it represents an indirect measurement of cell volume. We assumed that fluorescence was proportional to cell volume, which was validated using hypertonic solutions containing nonpermeating solutes. However, chemicals that can enter the cell have the potential to interact directly with calcein and influence its fluorescence. If CPAs affect calcein fluorescence, it may lead to inaccuracies in our permeability estimates. The potential for error due to interaction between calcein and CPA is examined in detail in the Supplementary Material Section S2 (Figures S4-S9), demonstrating that in most cases the permeability values reported here are expected to be accurate.

## 4. Conclusion

We have developed a high throughput method for assessing cell membrane permeability that is ~100 times faster than conventional techniques, while also enabling estimation of CPA toxicity using the same 96-well plate. This approach provides an efficient means of screening multiple compounds to identify chemicals with promising permeability and toxicity properties. We tested 5 common CPAs and 22 less common chemicals at both 4 °C and room temperature, with 23 passing the screening process. Overall, the methodology presented here holds great potential as an initial screening tool for identifying novel CPAs for advancing the field of cryopreservation, particularly in the context of preserving complex structures like organs and tissue constructs.

## 5. Materials and methods

### 5.1. Overview

Figure 1 illustrates the experimental method for high throughput measurement of cell membrane permeability and CPA toxicity. Cells cultured in a 96-well plates were first stained using calcein, a fluorescent dye that is confined to the cell cytoplasm. In the next step, the well plate was placed into a plate reader, which enabled automatic injection of CPA solutions into the wells while monitoring the fluorescence changes resulting from water and CPA transport through the cell membrane. Monitored fluorescent changes were used to determine the cell membrane permeability to water and CPA. Finally, the same plate was used to measure CPA toxicity. Toxic chemicals cause cell death and release calcein into the surrounding solution. Therefore, fluorescence was measured before and after removal of the solution from the wells, and the ratio between these fluorescence values was used to estimate the toxicity of the CPA.

### 5.2. Materials

Figure S1 shows the chemical structures of the candidate CPAs tested in this study. These chemicals were purchased from the following vendors: 1,3-propanediol (VWR), N,N-dimethylethanolamine (Fisher Scientific Company), 2-methoxyethanol (Thermo Scientific Chemicals), triethanolamine (Chem Impex International), diethylene glycol (VWR), 2,3-butanediol (MilliporeSigma), 2-methyl-1,3-propanediol (Fisher Scientific), N-methylacetamide (VWR), acetamide (Thermo Scientific Chemicals), 1,3-dihydroxyacetone (VWR), ethanolamine (MilliporeSigma), glycerol (Macron Fine Chemicals), DMSO (Fisher Chemical), propylene glycol (VWR), ethylene glycol (Macron Fine Chemicals), formamide (Sigma-Aldrich), sucrose (Macron Fine Chemicals). Several chemicals were used for making HEPES buffered saline (HBS) and phosphate buffered saline (PBS): MgCl_2_⋅6H_2_O (VWR), CaCl_2_⋅2H_2_O (Fisher Chemical), NaCl (VWR), KCl (EMD Millipore), HEPES (VWR), Na_2_HPO_4_ (Macron Fine Chemicals™), KH_2_PO_4_ (ChemProducts), HCl (MilliporeSigma), NaOH (VWR). Calcein-AM was purchased from Life Technologies. Finally, Dulbecco’s Modified Eagle Medium (Gibco), fetal bovine serum (Gibco), and Penicillin-Streptomycin (Gibco) were used to prepare cell culture media.

### 5.3. Test Solutions

Isotonic HBS was prepared as described previously [24]. The osmolality of the buffer was determined using an Advanced Micro Osmometer Model 3300 (Advanced Instruments, Norwood, MA) and was found to be within 2% of 300 mOsm/kg. To create hypo- and hypertonic solutions, distilled water or sucrose were added to the isotonic HBS. CPA was added to the isotonic HBS to prepare CPA solutions.

### 5.4. Cell culture

Bovine pulmonary artery endothelial cells (BPAEC, Cell Applications, Inc. in San Diego, CA) were cultured as described previously [25]. Cells were obtained from the vendor at passage 2 and subsequently expanded to passage 5. At this stage, they were cryopreserved. For each experiment, a vial of cryopreserved cells was thawed and seeded into a T-75 flask containing 15 mL of culture medium. Cells were then cultured for 24 to 30 hours, reaching approximately 80% confluency. Subsequently, they were harvested from the T-75 flask and seeded into clear-bottom black 96-well plates (Greiner Bio-One, Monroe, NC) at a density of 10,000 cells per well. The well plates were then cultured for approximately 2 days, at which point the cells had achieved ~80% confluency.

### 5.5. Calcein staining

To stain the cells, the wells were first washed with 200 μL of Dulbecco’s PBS containing calcium and magnesium. Subsequently, the cells were exposed to concentration of 1.67 μg/mL calcein-AM in PBS for a duration of 30 minutes within an incubator at 37 °C. To eliminate any remaining calcein-AM, the plates were washed twice with 200 μL of isotonic HBS. Finally, for the experiment, 100 μL of isotonic HBS was retained within the wells.

### 5.6. Quantifying cell membrane permeability

Cell membrane permeability was quantified using a FlexStation 3 plate reader (Molecular Devices), which is equipped with an automated system that enables immediate fluorescence measurement after fluid transfer for kinetic assays. This system seamlessly transfers liquids between the “source” plate and the “assay” plate using an 8-channel head capable of dispensing into an entire column of the well plate simultaneously. The system also automates analysis of the whole well plate by acquiring fluorescence data one column at time. In this experimental configuration, the assay plate contains cells stained with calcein, while the source plate holds CPA solutions and other complementary substances intended for introduction into the cells.

Before commencing the experiments, the assay and source plates were subjected to at least 15-minute incubation at either room temperature (25 ± 2 °C) or 4 °C, depending on the designated test temperature. This was done to ensure the plates achieved thermal equilibrium at the test temperature. To ensure consistency, the plate reader was adjusted to the test temperature at least one hour prior to the test.

An excitation wavelength of 480 nm and emission wavelength of 515-530 nm were employed to detect the fluorescence intensity. Fluorescence data were collected for 30 seconds while the cells were in the isotonic HBS. At the 30-second mark, 100 μL of the test solution was added to the well plate containing the cells, and fluorescence data collection was performed for 2 minutes at 1.5-second intervals. No cell losses due to fluid shear were observed after dispensing the test solution, as confirmed by microscopic images of the cells taken afterward. The plate reader settings resulted in injection of test solution at a rate of 100 μL/sec at the surface of the initial solution in the well. The resulting fluorescence data was used to estimate the cell membrane permeability to CPA and water.

To analyze the results, we normalized the data based on the fluorescence during exposure to isotonic HBS. First, we focused on the initial 30 seconds of the experiment, during which the cells were exposed solely to the HBS buffer. We calculated the average fluorescence during this period for each well and used this average fluorescence to normalize all the fluorescence measurements for that well. This normalization accounts for well-to-well differences in fluorescence, resulting in a normalized fluorescence during the initial isotonic period of ~1.0 for each well. An additional normalization was also performed based on measurements for the isotonic HBS control wells. For these control wells, 100 μL of isotonic HBS was dispensed at the 30 second mark, rather than CPA solution. Each well plate had 12-24 isotonic HBS control wells. Interestingly, we noticed a slight decrease in fluorescence when we added additional HBS buffer. To account for the potential effects of solution injection, we combined the data from the isotonic HBS wells to create an average fluorescence at each time point, and used this to normalize all of the fluorescence data for the well plate. After completing both normalization steps, we refer to the resulting normalized cell fluorescence as *F̄*. Figure S2 shows representative results illustrating the normalization process. For the isotonic HBS control wells, the resulting normalized fluorescence deviated by no more than 2% from the expected value of *F̄* = 1.0.

To enhance measurement accuracy, we chose to exclude wells in the plates with notably low fluorescence intensity, specifically those registering less than 400. Additionally, for a few wells in some of the 96-well plates at 4 °C, we observed fluid transfer errors, which were confirmed by measuring the amount of solution in each well. Therefore, we have eliminated the data related to these errors from our dataset.

The normalized cell fluorescence *F̄* was used to estimate the water and CPA permeability parameters using an approach similar to that described previously [24]. The two-parameter model was used to predict changes in cell volume due to transport of water and CPA across the cell membrane:

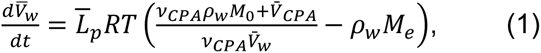

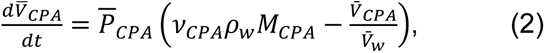

where *L̄_p_* = *L_p_*(*A*/*V_w_*_0_) is the effective water permeability, *P̄_CPA_* = *P_CPA_*(*A*/*V_w_*_0_) is the effective CPA permeability, A is the cell membrane surface area, *V_w_*_0_ is the cell water volume under isotonic conditions, *V̄_w_* = *V_w_*/*V_w_*_0_ is the normalized intracellular water volume, *V̄_CPA_* = *V_CPA_*/*V_w_*_0_ is the normalized intracellular CPA volume, R is the ideal gas constant, T is the absolute temperature, *ν_CPA_* is the molar volume of CPA, *ρ_w_* is the density of water (assumed to be 1 kg/L), *M*_0_ is the isotonic osmolality, *M*_e_ is the total extracellular osmolality, and *M_CPA_* is the extracellular CPA osmolality. The two-parameter model (Eqs. 1 and 2) was solved to determine the osmotically active cell volume,

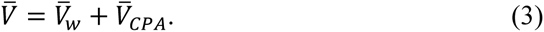

We assumed a linear relationship between normalized cell fluorescence *F̄* and the normalized osmotically active cell volume *V̄*,

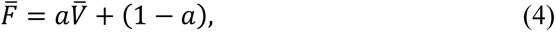

where *a* is a proportionality constant. For each plate, several wells were allocated to estimate the proportionality constant between *F̄* and *V̄*. These wells were exposed to hypertonic solution containing sucrose, which is considered a nonpermeating solute due to its extremely low membrane permeability. The extent of cell water volume loss after equilibrating with the hypertonic sucrose solution can be estimated using the Boyle-van ’t Hoff relationship,

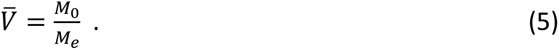

For each well plate, the proportionality constant *a* was first estimated from the sucrose wells. Then the best-fit permeability parameters *L̄_p_* and *P̄_CPA_* were estimated for each well by minimizing the sum of the error squared between the predicted and measured *F̄*-values. To obtain predicted *F̄*-values, *V̄* was determined using the two-parameter model and converted to *F̄* using the proportionality constant *a*. For experiments performed at room temperature, only the first 15 seconds after CPA exposure were used to determine the best-fit permeability values. For experiments at 4 °C, the first 90 seconds after CPA exposure were used.

The temperature dependence of water and CPA permeability can be assessed using the Arrhenius equation:

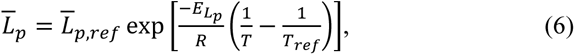

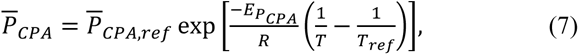

where 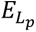 and 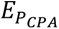 are the activation energy for water and CPA transport, *L̄_p,ref_* and *P̄_CPA,ref_* are the effective water and CPA permeability at a reference temperature *T_ref_*. The values of 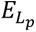 and 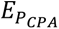 were derived through linear regressions applied to Arrhenius plots of the logarithmically transformed permeability parameters.

### 5.7. Quantifying CPA toxicity

For cell viability assessment, we utilized the same plate employed for cell membrane permeability testing. Following approximately 20 minutes of CPA exposure, we measured the fluorescence. Subsequently, we swiftly removed the liquid and conducted another fluorescence measurement. As depicted in Figure 1, cell death prompts the release of fluorescent dye into the surrounding solution. Therefore, extracting the solution from the wells leads to a notable reduction in fluorescence intensity within wells exposed to toxic CPAs. Cell viability was estimated using the ratio of fluorescence before and after removal of the liquid from the well. Because of the sequential injection of CPA solutions into different columns in the well plate during membrane permeability measurement, each column was exposed to CPA for a different amount of time. Viability analysis was only performed on wells that had been exposed to CPA for less than 35 minutes. At 4 °C, the range of CPA exposure times was 13 min to 34 min, resulting in an average exposure time of about 25 min. At room temperature, the range of exposure times was 8 min to 34 min, resulting in an average of about 20 min.

As shown in Figures S4, S7, and S10, exposure to some CPAs caused a downward trend in fluorescence, even before removing the liquid from the well to assess cell viability. This downward trend has the potential to confound viability estimation because of the resulting decrease in fluorescence during the time required to remove the liquid from the wells and make the second fluorescence measurement. To account for this, we fit an exponential decay model to the normalized fluorescence measured before removal of liquid from the well. Fluorescence data for different CPA exposure times was obtained from different columns in the plate. The best-fit exponential decay model was then used to estimate the expected value of normalized fluorescence corresponding to the time at which fluorescence was measured after CPA removal. Viability was determined from the ratio of these two values. This is illustrated for various CPAs in Figure S10.

### 5.8. Statistical analysis

The permeability data was found to have unequal variances for different CPAs, so the data was analyzed using Welch’s ANOVA, followed by Games-Howell tests for pairwise comparisons. The membrane permeability parameters for water and CPA were subjected to separate analyses. Significance was defined at a 95% confidence level. The entire analysis was conducted using Statgraphics 19 statistical software.

## Acknowledgements

This work was supported by funding from the National Institutes of Health (R01 EB027203). We would also like to acknowledge a generous donation to the laboratory of AZH from the Cryonics Institute.

