## Supplementary Material for "High throughput method for simultaneous screening of membrane permeability and toxicity for discovery of new cryoprotective agents"

#### S.1. Chemicals structures of candidate CPAs

The chemical structures of the candidate CPAs tested in this study are shown in Figure S1.

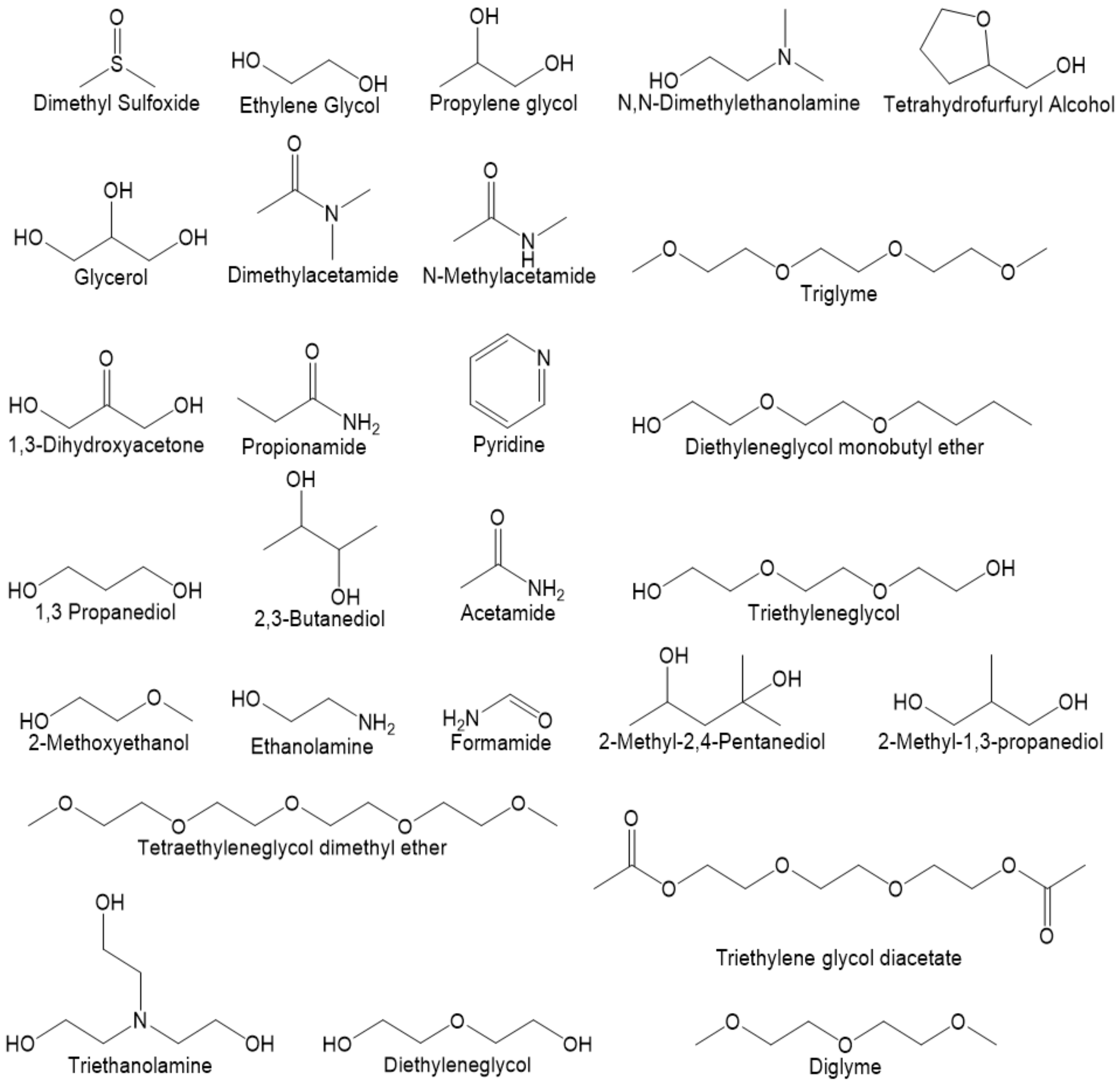


*Figure S1. Chemical structures of the candidate CPAs tested in this study.*

#### S.2. Fluorescence normalization for permeability measurement

The fluorescence data used for cell membrane permeability measurement was subjected to two normalization steps. Figure S2 shows normalized fluorescence values for the isotonic control wells on a representative 96-well plate. These wells initially contained 100 µL of isotonic buffer, then an additional 100 µL of isotonic buffer was injected after 30 seconds. In the first normalization step, the data in each well was normalized to the average fluorescence in the first 30 seconds. The resulting normalized fluorescence is illustrated in the top panel of Figure S2. As expected, this normalization resulted in a fluorescence of ~1.0 during the first 30 seconds. However, injection of additional isotonic buffer at the 30 second time point caused a slight drop in fluorescence. The second normalization step was performed to account for the impact of solution injection on normalized fluorescence. The fluorescence at each time point was normalized a second time using the corresponding normalized fluorescence of the isotonic control wells. The effect of this second normalization is illustrated in the bottom panel of Figure S2. Our analysis demonstrates that the final normalized fluorescence deviations do not exceed 2% from the expected value of F = 1.0.

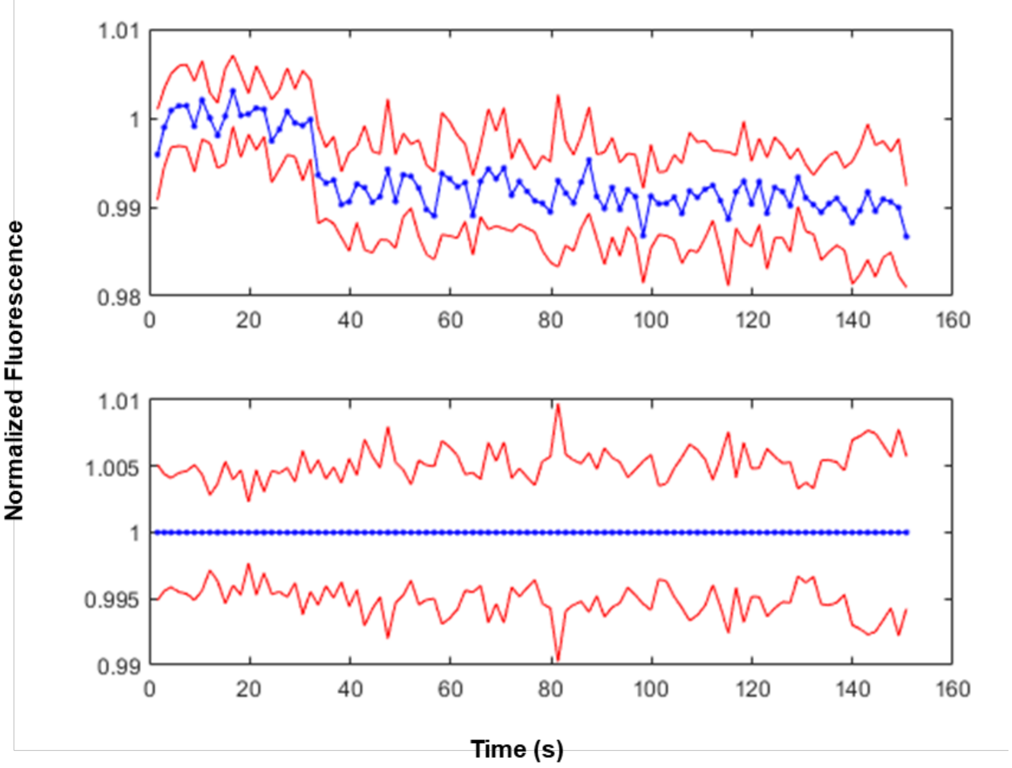


*Figure S2. Normalized fluorescence of the isotonic control wells for a representative 96-well plate. The top panel illustrates the first normalization step, and the bottom panel illustrates the second normalization step, showing the average normalized fluorescence (blue dots) ± the standard deviation (red lines).*

#### S.3. Factors affecting fluorescence after CPA exposure

Exposure to a solution containing permeating CPA causes cell volume loss and then gain. These volume changes can affect fluorescence by changing the environment around the intracellular calcein molecules. Previous studies indicate that calcein fluorescence is quenched by molecules present in the cell cytoplasm, and that cell volume changes affect fluorescence by altering the concentration of these quencher molecules [1]. To examine the relationship between cell volume and fluorescence, we exposed the cells to hypo- and hypertonic solutions containing nonpermeating solutes but lacking permeating CPAs. After reaching equilibrium in these solutions, the cell volume can be predicted from the Boyle-van 't Hoff relationship, enabling the measured fluorescence to be plotted against the predicted cell volume. As shown in Figure S3, a linear relationship between fluorescence and volume was observed under hypertonic conditions. However, the fluorescence under hypotonic conditions deviated from this linear trend. In this study, we used the linear fit to the hypertonic data in Figure S3 to convert between fluorescence and volume after CPA exposure.

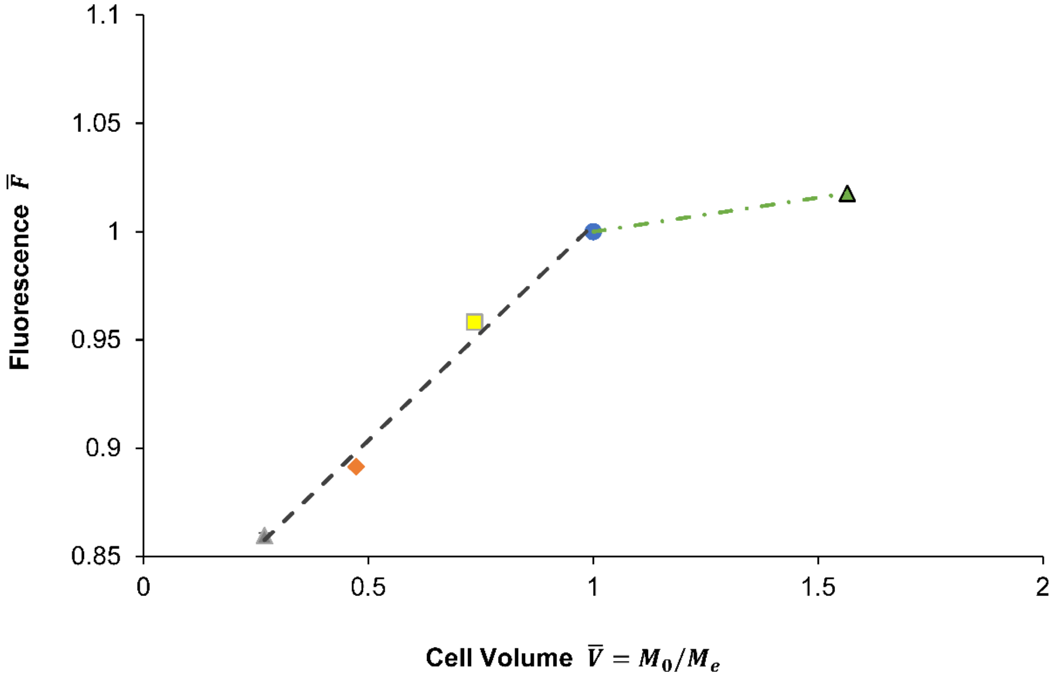


*Figure S3. Effect of cell volume on fluorescence for hypertonic and hypotonic solutions containing nonpermeating solutes. Each data point represents the average ± SEM of the equilibrium fluorescence for 10-12 wells. Experiments were performed at 25 °C. Error bars are shown but are small enough to make discerning difficult.*

Permeating CPA exposure also results in uptake of the CPA, which has the potential to affect fluorescence independent of changes in cell volume. When a CPA enters the cell, it could change the solvent environment around calcein and affect its fluorescence. The solvent environment around a fluorophore can affect its fluorescence by altering polarity or viscosity, or by direct interaction with the fluorophore (e.g., hydrogen bonding or complex formation) [2, 3]. We will refer to this as "solvent effects." Additionally, changes in the environment around calcein could affect the rate of bleaching or fading of fluorescence [2, 3]. To account for solvent effects and bleaching, and distinguish these mechanisms from fluorescence changes due to cell volume changes, we examined fluorescence data acquired approximately 10 to 50 minutes after CPA exposure. For these longer time points, the cells are expected to be in equilibrium with the CPA solution and no longer changing volume.

S.3.1. Analysis of equilibrium fluorescence after CPA exposure at 4 °C

Figure S4 shows the fluorescence after exposure to various candidate CPAs for up to 50 min at 4 °C. The data was acquired as a part of our normal experimental workflow for assessing CPA toxicity, which includes a fluorescence measurement for the whole well plate after CPA exposure. Each data point in Figure S4 represents a measurement for a single well. For all the permeating CPAs besides glycerol, we only included data points for wells exposed to the CPAs for long enough to be close to equilibrium, based on cell volume predictions using the two-parameter model (Eqs. 1-2). Glycerol had such a low permeability that none of the data points were predicted to be close to equilibrium. The data presented in Figure S4 allows us to examine the potential impact of bleaching and solvent effects under conditions where cell volume is not changing.

The presence of a downward slope over time in Figure S4 indicates bleaching. To assess bleaching, regression analysis was performed. The slope was found to be significant in 6 cases. For the remaining CPAs, the slope was not statistically different from zero, indicating negligible bleaching. For the 6 cases where bleaching was non-negligible, an exponential decay model was fitted to the data to determine a bleaching rate constant. The resulting rate constant $k$ is shown in Figure S3.

The data in Figure S4 can also be used to examine the impact of solvent effects. The equilibrium fluorescence after exposure to permeating CPAs is expected to be approximately 1.01 if cell volume is the only factor affecting fluorescence (see Figure S3). Solvent effects may cause deviation from this fluorescence value. To isolate solvent effects, we estimated the equilibrium fluorescence after correcting for bleaching. Three different approaches were used to estimate the equilibrium fluorescence. For CPAs that caused cells to reach equilibrium within the first 1.5 min, the equilibrium fluorescence was estimated using the last data points displayed in Figure 3 of the main manuscript. We assumed that bleaching was negligible over this short period. For CPAs in Figure S4 that resulted in negligible bleaching, the equilibrium fluorescence was calculated by taking the average over all the wells in the plate. For CPAs demonstrating bleaching in Figure S4, exponential extrapolation to zero was employed to estimate the equilibrium fluorescence.

The resulting equilibrium fluorescence results are shown in Figure S5. For most CPAs, the equilibrium fluorescence was close to 1.01, suggesting that solvent effects are negligible for most CPAs. However, some CPAs yielded an equilibrium fluorescence that deviated from 1.01 (e.g., acetamide, triethylene glycol), suggesting that solvent effects may be non-negligible in these cases. It is also noteworthy that exposure to the nonpermeating solute sucrose, which causes cell shrinkage, resulted in an equilibrium fluorescence that is substantially lower than the equilibrium fluorescence for all the CPAs. This suggests that cell volume is the dominant factor affecting fluorescence.

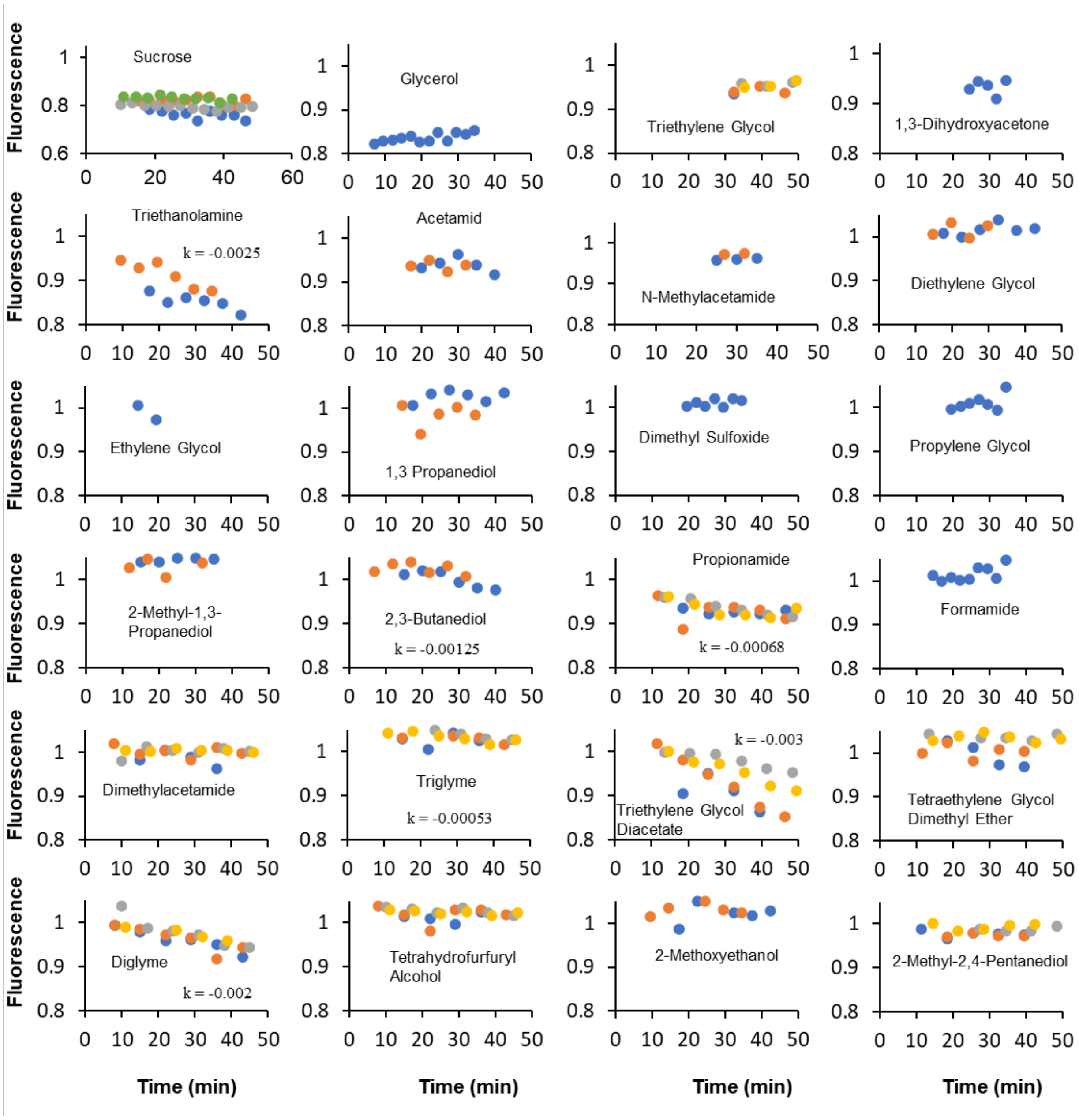


*Figure S4. Fluorescence after exposure to various candidate CPAs at 4 °C. The solutions had a total osmolality of 1000 ± 100 mOsm/kg. Different colors represent results for different well plates. At least one plate was studied for each CPA. In some cases, an exponential model was fitted to the data, resulting in the rate constant k (min$^{-1}$) shown on the graph.*

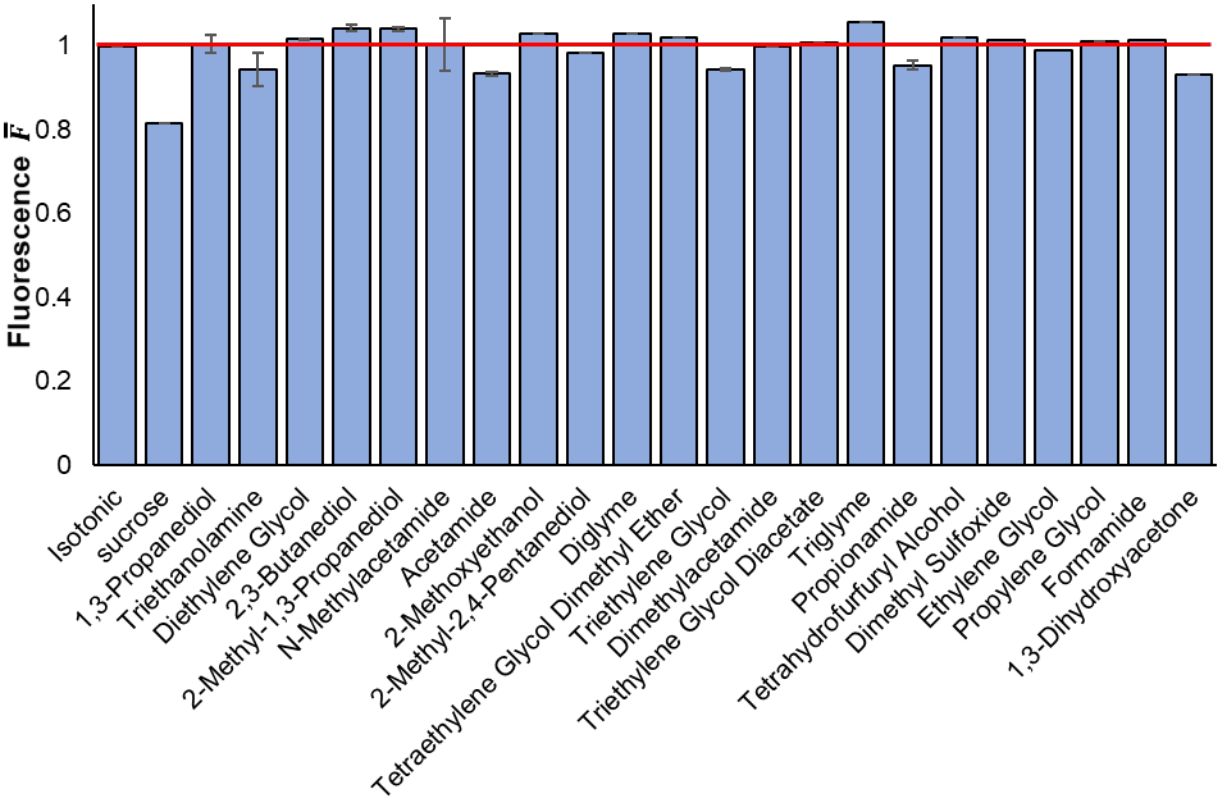


*Figure S5. Equilibrium fluorescence in the presence of candidate CPAs at 4 °C. The red line shows the expected equilibrium fluorescence value assuming cell volume is the only factor affecting fluorescence.*

The above analysis shows that in some cases, solvent effects and bleaching may have non-negligible effects on fluorescence. However, in the main manuscript, we assumed that these factors were negligible for fitting the data to estimate the cell membrane permeability to water and CPA. To examine potential errors caused by non-negligible bleaching and solvent effects, we re-examined the data using a mathematical model accounting for these factors:

$$\bar{F} = [e^{-kt}]\left[1 + \alpha\frac{N_{CPA}}{N_{CPA,equil}}\right][a\bar{V} + (1 - a)],$$

where α is a solvent effects constant (defined as $\bar{F}_{equil}/1.01 - 1$), $N_{CPA}$ is the moles of intracellular CPA, and the subscript "equil" denotes equilibrium conditions. In the equation above, the first term in brackets accounts for bleaching, the second term accounts for solvent effects, and the third term is identical to Eq. 4 from the main manuscript and accounts for the effects of cell volume changes on fluorescence.

Four different fits were performed, including the original fit, and fits accounting for solvent effects, bleaching, and a combination of solvent effects and bleaching. Figure S6 (top) shows the water permeability values for each of these four fits. Bleaching and solvent effects had a negligible effect on the best-fit water permeability for all CPAs except propionamide. This suggests that the original fit is accurate in these cases. For propionamide, accounting for bleaching and solvent effects reduced the best-fit water permeability by 2x, suggesting that the original fit may have slightly overestimated the water permeability.

Figure S6 (bottom) shows the CPA permeability values for each of the four fits. For all the CPAs except propionamide and triethylene glycol, the original CPA permeability values were within a factor of two of the best-fit values that accounted for bleaching and solvent effects. This indicates that the original CPA permeability values are reasonably accurate for most CPAs. For propionamide and triethylene glycol, accounting for bleaching and solvent effects increased the best-fit CPA permeability by 5x and 4x, respectively, suggesting that the original fit may have underestimated the CPA permeability in these cases.

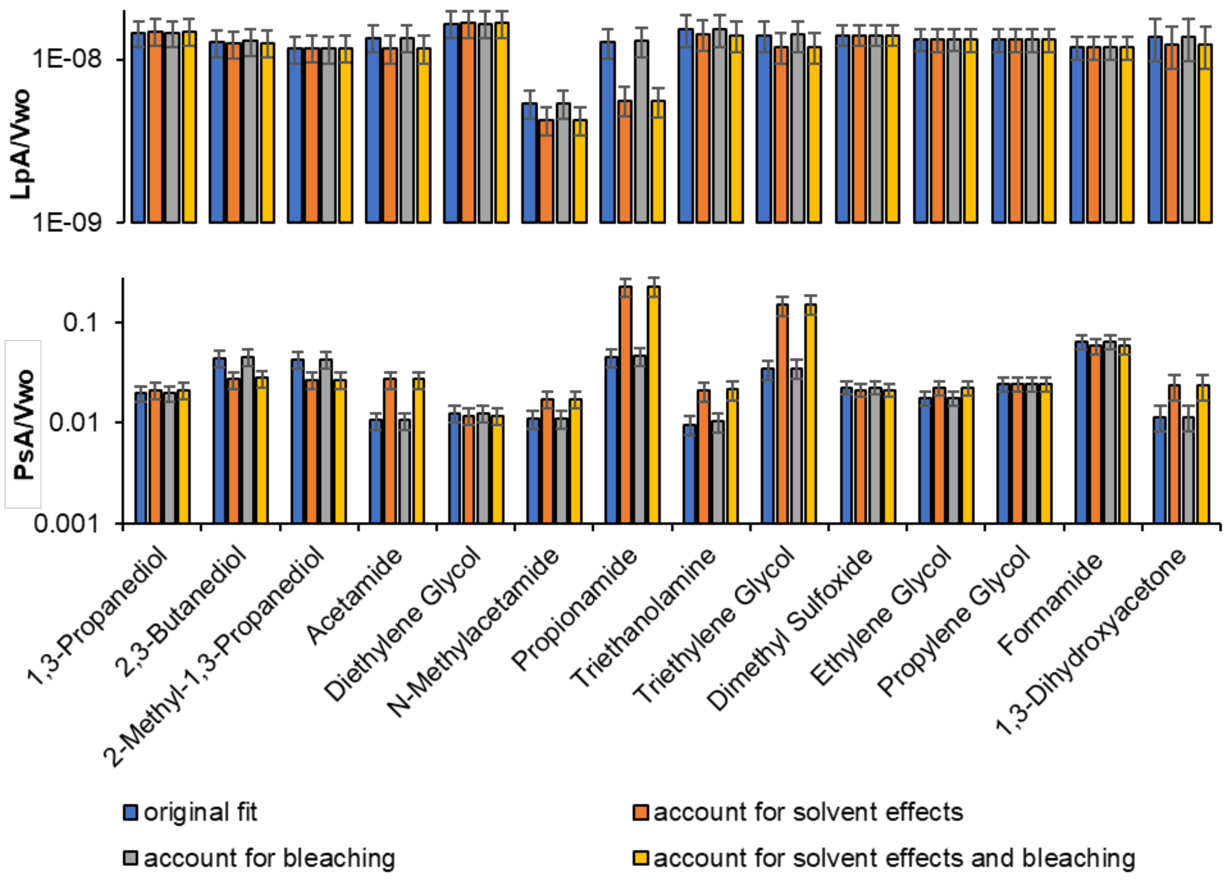


*Figure S6. Water and CPA permeability for different CPAs at 4 °C using various fitting methods, including fit without accounting for solvent effects and bleaching (i.e., original fit), fit accounting for solvent effects, fit accounting for bleaching, and fit accounting for both solvent effects and bleaching. Each data point represents the average ± SEM of a minimum of 13 replicates conducted across at least 3 plates.*

*S.3.2. Analysis of equilibrium fluorescence after CPA exposure at 25 °C.*

As shown in the Figure S7, the effect of bleaching at 25°C is greater than at 4°C. Overall, 15 chemicals exhibited a downward slope that was statistically significant based on regression analysis (compared to 6 at 4 °C). In addition, the bleaching rate constants were higher, indicating faster bleaching.

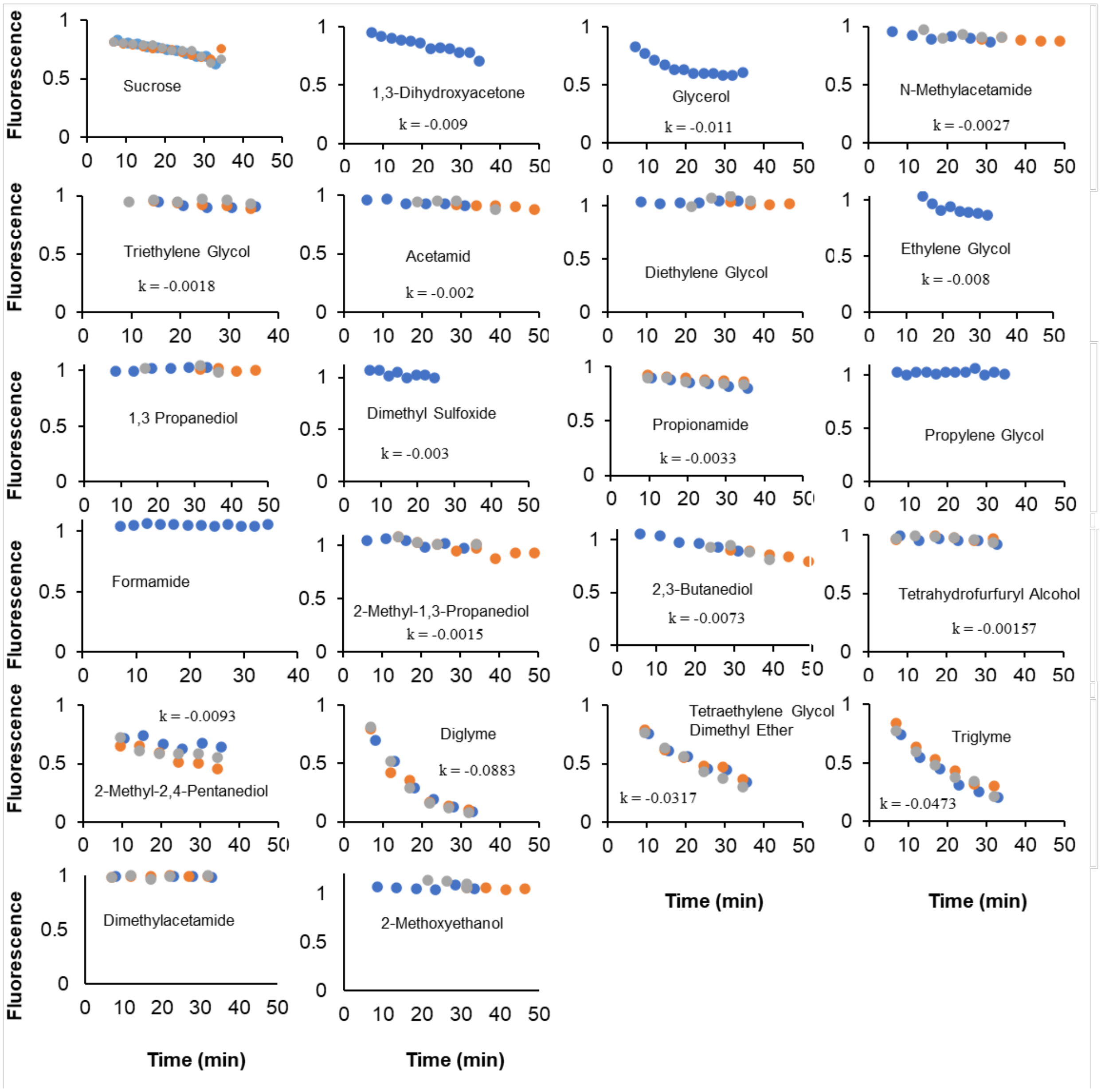


*Figure S7. Fluorescence after exposure to various candidate CPAs at 25 °C. The solutions had a total osmolality of 2000 ± 100 mOsm/kg, except for sucrose, which had an osmolality of 1000 ± 100 mOsm/kg. Different colors represent results for different well plates. At least one plate was studied for each CPA. In some cases, an exponential model was fitted to the data, resulting in the rate constant k (min$^{-1}$) shown on the graph (see text for details).*

Figure S8 shows the equilibrium fluorescence values estimated using the data in Figure S7. Most equilibrium fluorescence values are within the expected range, suggesting negligible solvent effects. However, 1,3-dihydroxyacetone, propionamide, and glycerol had relatively low equilibrium fluorescence values, possibly due to solvent effects. Note that for glycerol there is significant uncertainty in the estimate for the equilibrium fluorescence shown in Figure S7. The data is from a single well plate, and the exponential fit used to correct for bleaching was not very accurate ($R^2 = 0.80$).

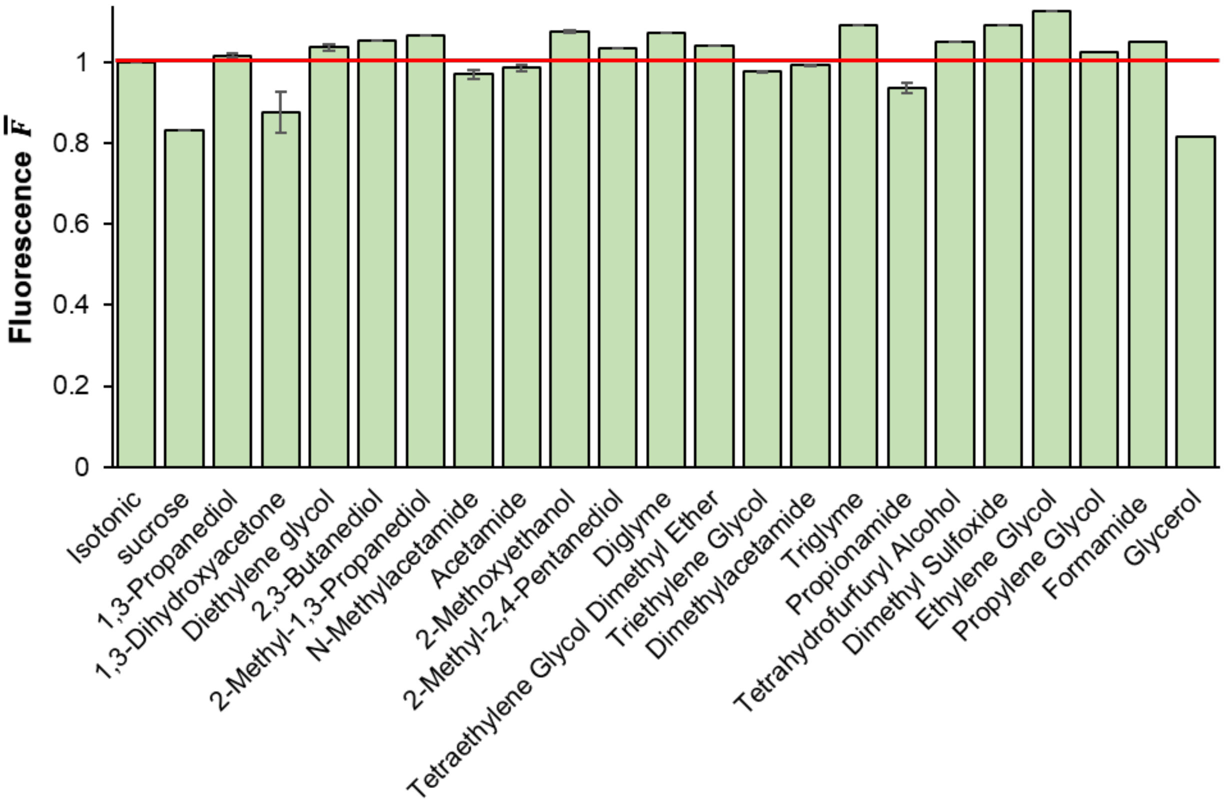


*Figure S8. Equilibrium fluorescence in the presence of candidate CPAs at 25 °C. The red line shows the expected equilibrium fluorescence value assuming cell volume is the only factor affecting fluorescence.*

Figure S9 presents best-fit cell membrane permeability values using our original fluorescence model, as well as fluorescence models augmented to account for bleaching and/or solvent effects. Similar to the results at 4 °C, most CPAs yielded cell membrane permeability values within the same range for all fits at 25 °C. Thus, for most chemicals, our original permeability fits are expected to be reasonably accurate. However, the variation at 25 °C is higher than at 4 °C. More specifically, propionamide and glycerol show substantial differences in water permeability, while propionamide, glycerol, formamide, and triethylene glycol show substantial differences in CPA permeability (Figure S9). For these cases, solvent effects and/or bleaching may have led to error in the permeability values reported in the main manuscript.

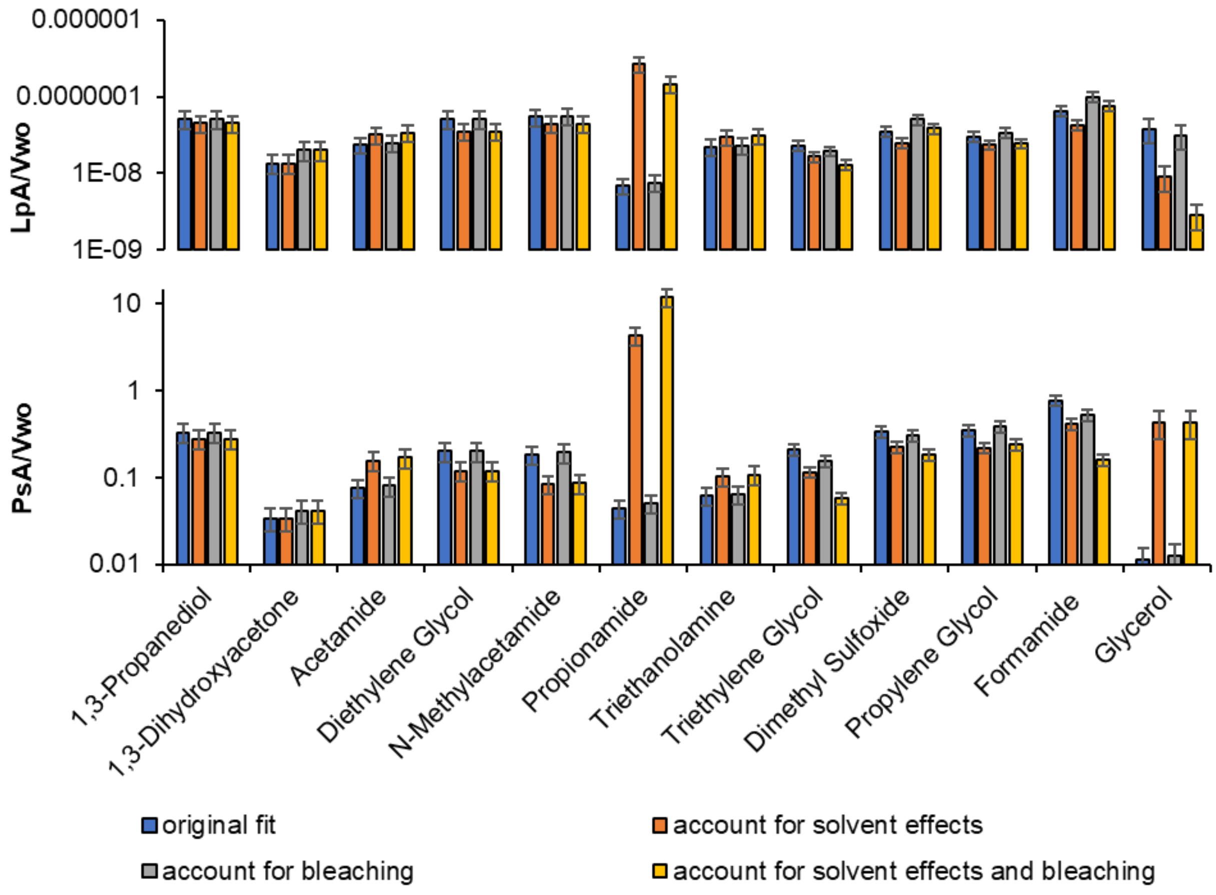


*Figure S9. Water and CPA permeability for different CPAs at 25 °C using various fitting methods, including fit without accounting for solvent effects and bleaching (i.e., original fit), fit accounting for solvent effects, fit accounting for bleaching, and fit accounting for both solvent effects and bleaching. Each data point represents the average ± SEM of a minimum of 13 replicates conducted across at least 3 plates.*

### S.4. Analysis of fluorescence for assessing CPA toxicity

To assess loss of cell viability caused by CPA exposure, fluorescence was measured before adding the CPA, after ~20 min CPA exposure, and after removing the CPA solution from the wells. The fluorescence after CPA exposure and after removing the solution was normalized using the initial fluorescence before CPA exposure. The resulting normalized fluorescence values are plotted against CPA exposure time in Figure S10 for four different CPAs. For some CPAs, the fluorescence after CPA exposure decreased as exposure time increased. To account for this, the data was fitted using an exponential decay model (red lines). Viability was estimated from the ratio of the fluorescence after removal of the solution (green circles) to the fluorescence predicted using the exponential decay model at the same time point. The top left panel illustrates a nontoxic CPA that did not exhibit a notable decrease in fluorescence with time. The top right panel shows a nontoxic

CPA that exhibited substantial decrease in fluorescence with time. The bottom panels show representative results for toxic CPAs.

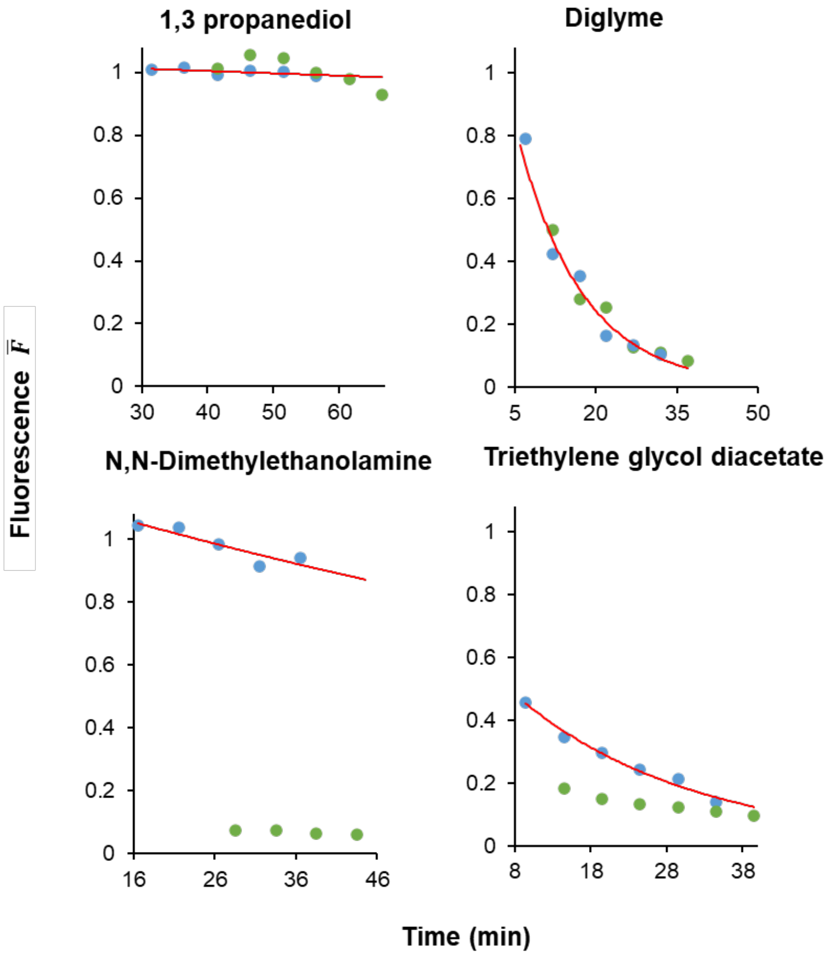


*Figure S10.Analysis of fluorescence data for viability measurement for four representative CPAs. If green dots are below the red line, it is a sign of CPA toxicity (blue dots: after exposure but before the removal of the solution from the wells, green dots: after removal of the solution, red line: exponential decay fit to blue dots).*

### S.5. Comparison with similar studies

To compare the data generated by our new method with other studies, we conducted a comparative analysis with two different studies, as illustrated in Figure S11 and S12.

First we aimed to assess the alignment between our new high-throughput method and our previous study [4], which also investigated the membrane permeability of bovine pulmonary artery endothelial cells. As depicted in Figure S11, a linear relationship was observed between our new solute permeability data and our previous data for both 4 °C and room temperature. Both graphs exhibit high $R^2$ values (0.95 at 4 °C and 0.91 at room temperature). The slope of the linear fit revealed that our method yielded approximately 2- and 4-times higher permeability values than our previous data for 4 °C and room temperature, respectively.

Next, we compared our permeability data to published data for human red blood cells [5]. Figure S12 demonstrates an approximately linear trend between our measured data and this study at both 4 °C and room temperature, with $R^2$ values of 0.80 and 0.71, respectively. Additionally, for solutes categorized as fast-permeable in our study, the corresponding permeability in red blood cells was also high.

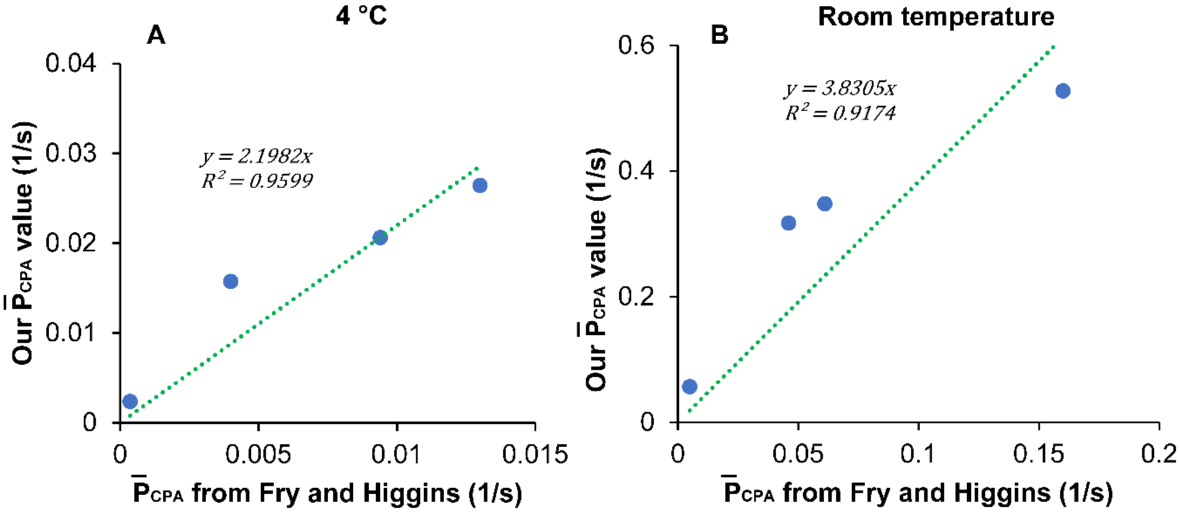


*Figure S11. Comparison of our solute permeability data with the permeability data from Fry and Higgins [4] at 4 °C and room temperature. Solute permeability values are shown for the following solutes: glycerol, DMSO, propylene glycol, and ethylene glycol.*

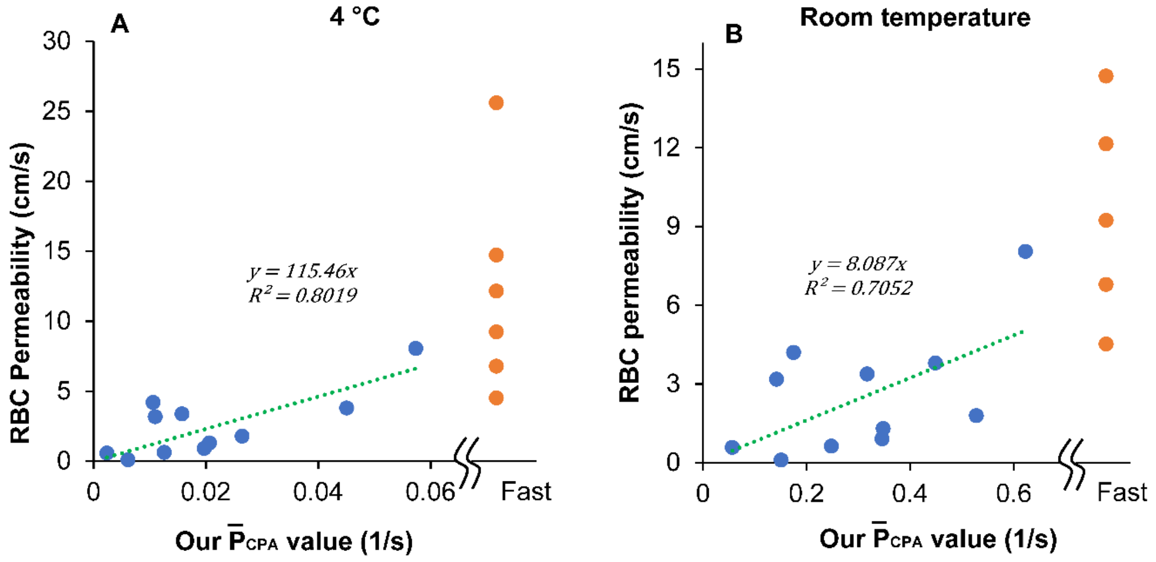


*Figure S22. Comparison of our solute permeability data with the permeability of human red blood cells [5] A) Comparison to our data at 4°C. B) Comparison to our data at 25°C. Solute permeability values are shown for the following solutes: glycerol, triethylene glycol, acetamide, methyl acetamide, diethylene glycol, ethylene glycol, 1,3-propanediol, dimethyl sulphoxide, 1,2-propanediol, propionamide, formamide. ethylene glycol monomethyl ether (i.e., 2 methoxyethanol), dimethyl acetamide, tetrahydrofurfuryl alcohol, 2-methyl-2,4-pentanediol, tetraethylene glycol dimethyl ether, triethylene glycol diacetate.*
